# Modeling interregional propagation of α-synuclein in human striatal–midbrain assembloids

**DOI:** 10.64898/2026.05.11.723670

**Authors:** Kaneyasu Nishimura, Naoya Amimoto, Yuya Sato, Mieko Morishima, Mamoru Sakaibara, Ayana Haratake, Mai Nakano, Naoko Kaneko, Hideaki Yamamoto, Ayumi Hirano-Iwata, Takashi Tanii, Jun Takahashi, Kazuyuki Takata, Yoshito Masamizu

**Author notes:** **Corresponding author:** Kaneyasu Nishimura, Ph.D., Laboratory of Functional Brain Circuit Construction, Graduate School of Brain Science, Doshisha University, Kyotanabe 610-0394, Japan.

## Abstract

Parkinson’s disease is characterized by the progressive degeneration of dopaminergic (DA) neurons in the substantia nigra that project to the striatum. Postmortem studies have found that α-synuclein pathology spreads across brain regions in a prion-like manner. Human three-dimensional neural tissues provide experimentally tractable platforms for clarifying the molecular pathology of synucleinopathy that overcome the limitations of postmortem brain studies. In this study, we established a human striatal–midbrain assembloid model enabling spatially and temporally controlled α-synuclein induction after formation of the nigrostriatal pathway within hStrMAs. Human induced pluripotent stem cells (hiPSCs) were differentiated into striatal spheroids and midbrain spheroids, which were fused to reconstruct the nigrostriatal pathway *in vitro*. Histological, electrophysiological, and single-cell RNA sequencing analyses confirmed the generation of region-specific neuronal populations, including striatal GABAergic neurons and midbrain DA neurons, and demonstrated that distinct forebrain and ventral midbrain identities were preserved within assembloids. To model synucleinopathy, we generated a doxycycline-inducible α-synuclein hiPSC line, permitting selective induction of α-synuclein expression in the striatal region after assembloid formation. Spatially restricted α-synuclein induction resulted in the progressive accumulation of phosphorylated and fibrillar α-synuclein in midbrain DA neurons, indicating interregional propagation through established neural circuits. Transcriptomic analysis of the midbrain region revealed significant downregulation of DA identity and synaptic function genes, suggesting functional impairment of DA neurons following α-synuclein propagation. These findings demonstrate that the human neural circuit architecture contributes to the interregional propagation of α-synuclein pathology. Our controllable human assembloid platform provides a powerful experimental framework for dissecting the mechanisms of pathological protein spread and developing therapeutic strategies targeting molecular pathology in neurodegenerative diseases.

**Significance statement:** One pathological feature of Parkinson’s disease is the progressive spread of α-synuclein pathology across brain regions. However, the molecular mechanisms underlying this process remain poorly understood. In this study, we established a human striatal–midbrain assembloid model in which α-synuclein expression can be spatially restricted and temporally induced after the nigrostriatal pathway is formed, thus enabling the experimental dissection of interregional α-synuclein propagation across brain regions. We demonstrated that α-synuclein pathology initiated in the striatal region propagated to midbrain dopaminergic neurons, leading to disease-relevant molecular and transcriptional alterations. These findings provide direct evidence that human neural circuits contribute to the spread of synucleinopathy and offer a controllable human model for studying disease progression and therapeutic strategies.

## Introduction

Parkinson’s disease (PD) is a progressive neurodegenerative disorder characterized by the loss of dopaminergic (DA) neurons in the substantia nigra pars compacta (SNpc) (1). DA neurons in the SNpc primarily project to the striatum, in which the axons of DA neurons project and input dopamine to the striatum, and impairment of the nigrostriatal pathway causes motor symptoms, including resting tremor, rigidity, and akinesia, in patients with PD (2). A key pathological feature of PD is the formation of Lewy bodies primarily composed of α-synuclein (3). In addition, missense mutations and duplication of the α-synuclein–encoding gene *SNCA* have been identified in familial PD, leading to increased α-synuclein expression and accumulation and subsequent DA neuron death in the SNpc (4–7). Another pathological feature is the progressive accumulation and propagation of α-synuclein across brain regions (8, 9). Postmortem studies have revealed that α-synuclein propagates across brain regions in a prion-like manner, and its distribution is correlated with the severity of motor symptoms during disease progression (10, 11). Although this concept provides a useful anatomical framework for understanding PD pathology, the cellular and circuit-level mechanisms underlying interregional α-synuclein propagation in the human brain remain incompletely understood. In rodent models, inoculation of α-synuclein preformed fibrils (PFFs) into the striatum induces the aggregation and spread of α-synuclein pathology across brain regions (12, 13). Two-dimensional (2D) neuronal cultures were used to further investigate cell-to-cell transmission and Lewy body-like inclusion formation (14–16). These *in vivo* and *in vitro* experimental platforms strongly contributed to clarification of the pathogenesis triggered by abnormal α-synuclein propagation.

Three-dimensional (3D) and self-organized neural organoids derived from human embryonic stem cells (hESCs) or induced pluripotent stem cells (hiPSCs) have emerged as promising platforms for modeling region-specific neural development and neurological disorders (17–19). More recently, neural assembloids generated by fusing distinct brain region-specific organoids have provided significant advantages for investigating interregional neural circuit formation and neural circuit-dependent pathological changes (20–25). Indeed, human striatal–midbrain assembloids (hStrMAs) can reconstruct the nigrostriatal pathway (26–29). Therefore, an experimental system that enables the induction of pathological changes of α-synuclein after nigrostriatal pathway establishment would permit the dissection of interregional α-synuclein propagation following neural circuit formation in assembloids.

Previously, we established efficient protocols for generating striatal and midbrain neurons using chemically defined medium containing small molecules that were replaced with protein components (30–32). Building on these previous protocols, we have established an hStrMA platform consisting of region-specific spheroids derived from hiPSCs, in which α-synuclein expression can be induced selectively in the striatal region after neural circuit formation. By combining a doxycycline (DOX)-inducible α-synuclein expression system with subsequent assembly of striatal and midbrain spheroids, this approach provides an experimental model for investigating the molecular pathology of α-synuclein after nigrostriatal pathway formation. Using this experimental platform, we directly examined whether α-synuclein pathology specifically initiated in the striatal region propagated to midbrain DA neurons. Our findings demonstrated that spatially restricted induction of α-synuclein in hStrMAs leads to the progressive accumulation of α-synuclein pathology in the midbrain region, providing experimental evidence that the neural circuit contributes to interregional α-synuclein propagation. This controllable human assembloid system provides a framework for dissecting the mechanisms of pathological protein spread and selective neuronal vulnerability in neurodegenerative diseases.

## Results

### Generation of human striatal spheroids (hStrSs) and human midbrain spheroids (hMSs) from hiPSCs

We previously established efficient xeno-free 2D culture protocols for generating striatal GABAergic and midbrain DA neurons from hESCs/hiPSCs by recapitulating key aspects of human brain development (**Fig. S1A–C**) (30–32). Briefly, neuronal derivation was achieved by dual SMAD inhibition using the bone morphogenic protein pathway inhibitor LDN193189 and transforming growth factor-β (TGF-β)/activin/nodal pathway inhibitor A83-01 in chemically defined Essential 6 medium. Specification into the lateral ganglionic eminence (LGE) was achieved using the WNT pathway inhibitor XAV939 and Smoothened receptor agonist purmorphamine (**Fig. S1B**). Meanwhile, caudal ventral midbrain (VM) identity was induced by purmorphamine, the glycogen synthase kinase 3β inhibitor CHIR99021, and fibroblast growth factor 8B (FGF8B) (**Fig. S1C**). At the neural progenitor stage, LGE neural progenitors expressed the neural progenitor marker SOX2, telencephalic marker FOXG1, forebrain–midbrain marker OTX2, and LGE marker GSH2 on day 17, whereas caudal VM neural progenitors expressed SOX2 and OTX2 alongside the VM markers LMX1, FOXA2, and EN1 on day 16 (**Fig. S1D**), indicating region-specific patterning consistent with striatal and caudal VM identities. Upon long-term maturation culture, striatal neural progenitors differentiated into mature neurons expressing the neuronal markers MAP2 and synapsin and formed extensive neural networks by day 75. These cells also expressed the striatal neuron markers DARPP32 and CTIP2 (**Fig. S1E**). In addition, a fraction of neurons in the striatal culture exhibited spontaneous calcium transient on day 99 (**Fig. S1F**). Similarly, caudal VM neural progenitors matured into midbrain DA neurons expressing tyrosine hydroxylase (TH), NURR1 (also known as NR4A2), and FOXA2 on day 75 (**Fig. S1G**).

Next, we adapted these induction protocols to 3D hiPSC cultures to generate hStrSs and hMSs. We first generated hStrSs using a 3D culture system according to the 2D protocol for striatal neuron induction (**Fig. 1A**). Immunocytochemistry demonstrated that SOX2-, FOXG1-, and GSH2-positive neural progenitors were predominantly induced by day 17 (**Figs. 1B and S2A**). These progenitors subsequently matured into striatal neurons expressing MAP2, synapsin, DARPP32, GAD65/67, and CTIP2 by day 75 (**Figs. 1C and S2B**). To characterize transcriptional changes during striatal differentiation and maturation, we performed bulk RNA sequencing (RNA-seq) to compare hStrSs on days 17 and 75. The neural progenitor markers *SOX1*, *SOX2*, *ASCL1*, and *NES* and the LGE progenitor markers *GSX2* and *DLX2* were markedly downregulated and the neuronal markers *SYN1*, *MAP2*, and *MAPT* and the striatal neuron-associated genes *GAD1*, *GAD2*, *SST*, and *ISL1* were upregulated on day 75 compared with their expression on day 17 (**Fig. 1D**). Gene Ontology (GO) analysis of genes upregulated on day 75 revealed significant enrichment of biological processes related to neuronal maturation, including ‘modulation of chemical synaptic transmission’ and ‘regulation of trans-synaptic signaling’ (**Fig. 1E**). In addition, enrichment of cellular component terms such as ‘synaptic membrane’ and ‘transporter complex’ was noted (**Fig. S2C**), whereas Kyoto Encyclopedia of Genes and Genomes (KEGG) pathway analysis highlighted significant enrichment of ‘neuroactive ligand–receptor interaction’ and ‘neuroactive ligand signaling’ (**Fig. S2D**). To further examine the regional identity of hStrSs, we mapped the bulk RNA-seq data obtained on day 75 onto 3D *in situ* hybridization datasets from the Allen Brain Atlas using the VoxHunt algorithm (33). Day 75 hStrSs exhibited the highest transcriptional similarity to the subpallial region of the E13.5 mouse brain (**Fig. 1F**). Finally, based on whole-cell patch–clamp recordings, striatal medium spiny neurons (MSNs) within hStrSs exhibited a late-spiking firing pattern, a hallmark electrophysiological feature of striatal MSNs (34), on day 152 (**Fig. S2E**).

**Figure 1.**
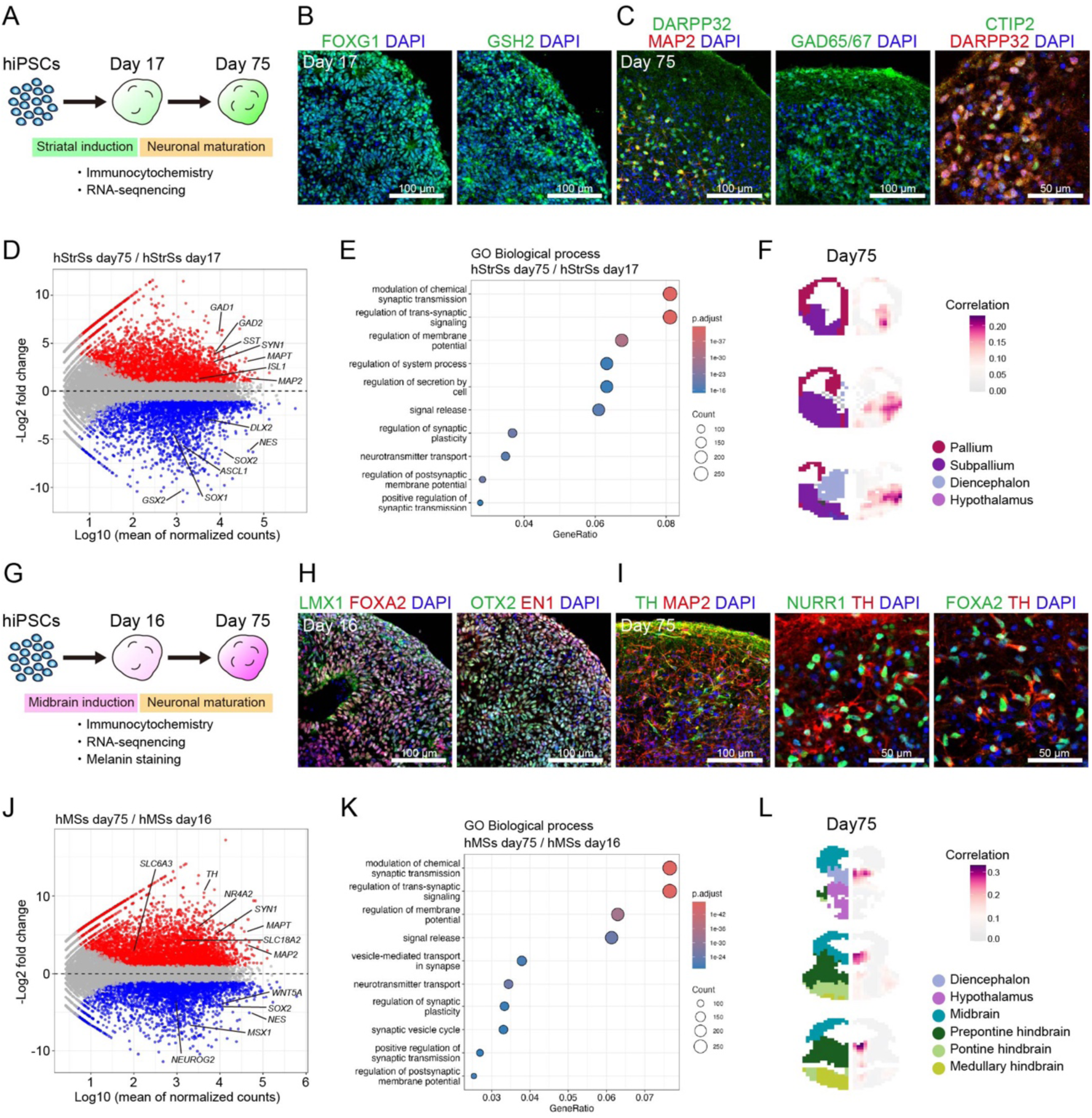
Induction and characterization of hStrSs and hMSs from hiPSCs. (**A**) Schematic illustration of the experimental procedure of hStrS generation from hiPSCs. (**B**) Immunostaining for FOXG1 and GSH2 on day 17. (**C**) Immunostaining for synapsin, MAP2, DARPP32, GAD65/67, and CTIP2 on day 75. (**D**) MA plot representing DEGs between days 17 and 75. Genes upregulated on day 75 are presented in red, and genes downregulated on day 75 are presented in blue (adjusted p-value < 0.05, |log2 fold change| > 1). (**E**) Top 10 upregulated GO terms on day 75 versus day 17. (**F**) VoxHunt spatial brain mapping of day 75 bulk RNA-seq data onto E13.5 Allen Developing Mouse brain Atlas data. (**G**) Schematic illustration of the experimental procedure of hMS generation from hiPSCs. (**H**) Immunostaining for LMX1, FOXA2, OTX2, and EN1 on day 16. (**I**) Immunostaining for MAP2, TH, NURR1, and FOXA2 on day 75. (**J**) MA plot representing DEGs between days 16 and 75. Genes upregulated on day 75 are presented in red, and genes downregulated on day 75 are presented in blue (adjusted p-value < 0.05, |log2 fold change| > 1). (**K**) Top 10 upregulated GO terms on day 75 versus day 16. (**L**) VoxHunt spatial brain mapping of day 75 bulk RNA-seq data onto E13.5 Allen Developing Mouse brain Atlas data.

Similarly, we generated hMSs using a 3D culture system based on the 2D protocol for midbrain DA neuron induction (**Fig. 1G**). Immunocytochemistry revealed the existence of SOX2-, LMX1-, FOXA2-, OTX2-, and EN1-positive cells by day 16 (**Figs. 1H and S1F**). These neural progenitors subsequently matured into midbrain DA neurons expressing MAP2, synapsin, TH, NURR1, and FOXA2 by day 75 (**Figs. 1I and S2G**). In addition, neuromelanin granules were detected on day 75 (**Fig. S2H**). RNA-seq indicated that the neural progenitor markers *SOX2* and *NES* and the VM progenitor markers *MSX1*, *NEUROG2*, and *WNT5A* were markedly downregulated and the neuronal markers *SYN1*, *MAP2*, and *MAPT*, and midbrain neuron-associated genes *TH*, *SLC18A2* (also known as *VMAT2*)*, SLC6A3* (also known as *DAT*), and *NR4A2* were upregulated on day 75 compared with their expression on day 16 (**Fig. 1J**). GO analysis of genes upregulated on day 75 revealed significant enrichment of biological processes related to neuronal maturation, including ‘modulation of chemical synaptic transmission’ and ‘regulation of trans-synaptic signaling’ (**Fig. 1K**), and upregulation of cellular component terms, such as included ‘synaptic membrane’ and ‘transporter complex’ (**Fig. S2I**). Meanwhile, KEGG pathway analysis highlighted significant enrichment of ‘neuroactive ligand–receptor interaction’ and ‘neuroactive ligand signaling’ (**Fig. S2J**). VoxHunt analysis demonstrated that day 75 hMSs displayed high transcriptional similarity to the midbrain region of the E13.5 mouse brain (**Fig. 1L**). According to whole-cell patch–clamp recordings, midbrain DA neurons within hMSs exhibited spontaneous pacemaking neuronal activity, a hallmark electrophysiological feature of DA neurons (35), on day 126 (**Fig. S2K**).

Next, we comprehensively validated the regional identities and differentiation states of hStrSs and hMSs by bulk RNA-seq at both the neural progenitor (hStrSs, day 17; hMSs, day 16) and mature neuronal stages (hStrSs, day 75; hMSs, day 75) **(Fig. 2A**). First, quantitative PCR confirmed appropriate region-specific marker expression at the neural progenitor stage, as *GSH2* was enriched in hStrSs and *LMX1A* and *FOXA2* were preferentially expressed in hMSs (**Fig. 2B**). Clustering analysis of bulk RNA-seq datasets demonstrated that neurospheres segregated according to brain regional identity and differentiation stage, indicating high intragroup similarity and clear intergroup separation (**Fig. S3A**). Principal component analysis (PCA) further supported this segregation, with samples clustering distinctly by regional identity and maturation status (**Fig. 2C**). The first principal component (PC1) primarily reflected neuronal differentiation and maturation-related gene expression changes. PC2 was associated with neural progenitor-related gene expression, and PC3 corresponded to region-specific transcriptional signatures distinguishing striatal and VM identities (**Fig. S3B**). Consistent with these findings, gene set enrichment analysis demonstrated that at both the neural progenitor and mature neuronal stages, LGE- and striatal-related genes were significantly enriched in hStrSs, and floor plate- and VM-related genes were enriched in hMSs (**Figs. 2D and S3C–D**). These data confirmed the successful generation of hStrSs and hMSs that closely resemble their corresponding brain regions *in vivo*.

**Figure 2.**
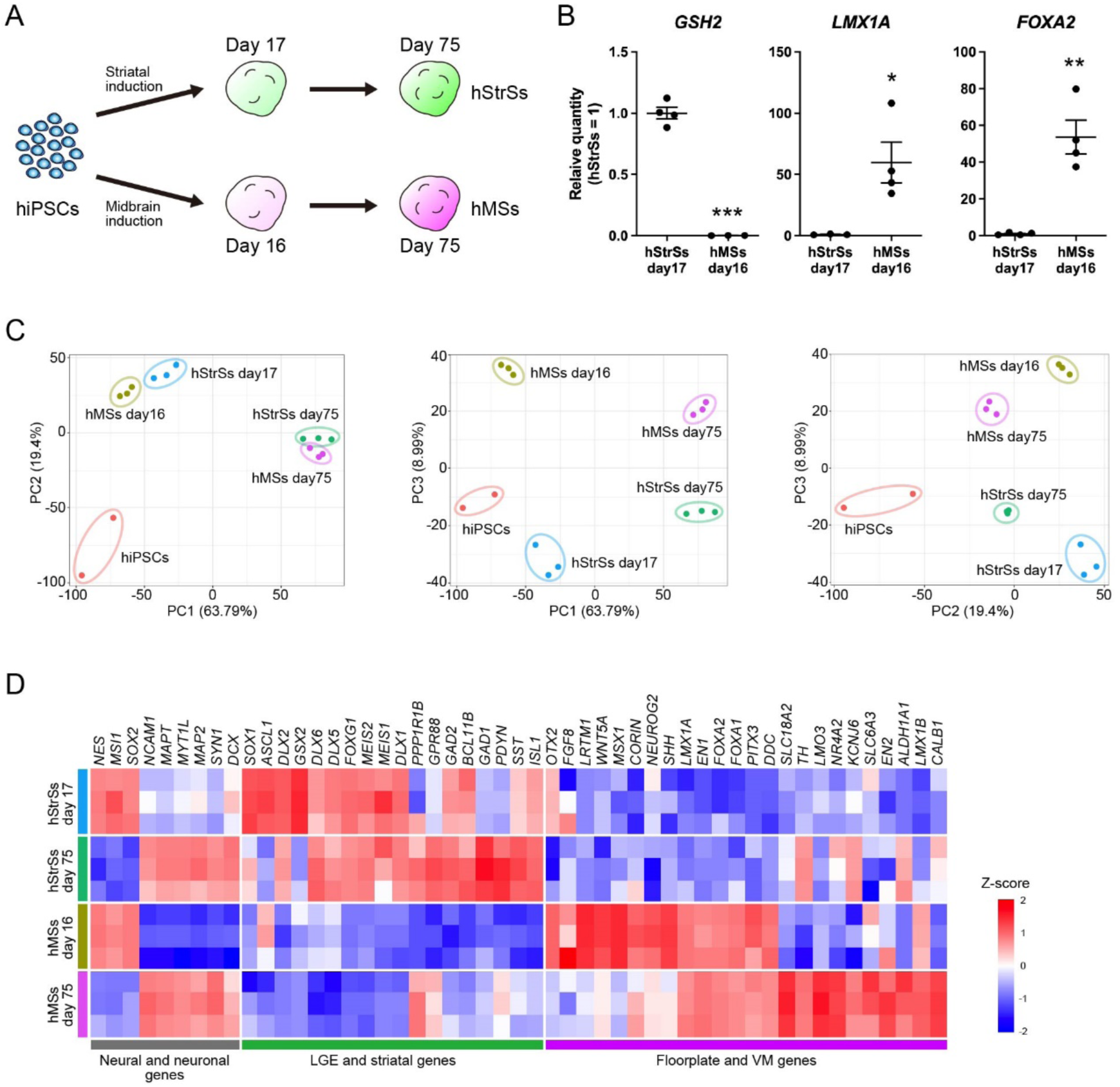
Comprehensive analysis of hStrSs and hMSs by bulk RNA-seq. (**A**) Schematic illustration of the experimental procedure for generating hStrSs and hMSs from hiPSCs. (**B**) Comparison of *GSH2*, *LMX1A*, and *FOXA2* expression between hStrSs on day 17 and hMSs on day 16. Significance (Student’s *t*-test): \**P* < 0.05; \*\**P* < 0.01; \*\*\**P* < 0.001 (n = 3–4, mean ± SD). (**C**) PCA of RNA-seq data from hiPSCs, hStrSs, and hMSs. (**D**) Heatmap presenting representative genes related to neural and neuronal genes, LGE and striatal genes, and floorplate and VM genes.

### Generation of hStrMAs

To generate hStrMAs, a single hStrS and hMS were cocultured in one well of ultralow-attachment U-bottom 96-well plate on day 30 of differentiation (**Figs. 3A and S4A, B**). To optimize the spheroid size, hStrSs and hMSs were initially differentiated from 12,000 and 6,000 cells per well, respectively. Under these conditions, the two spheroids rapidly fused to each other (**Figs. 3B–C and S4C–D; Movie S1**). Quantitative analysis revealed that the intersurface length between hStrSs and hMSs significantly increased by 5 days postfusion (dpf) compared with the findings immediately after fusion (**Fig. 3D, E**). Importantly, immunocytochemistry at 5 dpf confirmed that regional identities were preserved, with distinct striatal and midbrain domains maintained within the fused spheroids (**Fig. 3F**). By 30 dpf, hStrMAs predominantly consisted of MAP2-positive mature neurons, and the striatal and midbrain regions remained clearly segregated **(Fig. 3G**). The striatal region was enriched with GAD65/67-positive inhibitory neurons, whereas the midbrain region preferentially contained TH-positive DA neurons (**Fig. 3H**). Notably, TH-positive midbrain DA axons extended into the striatal region (**Fig. 3I**), and DA neural fibers were observed near DARPP32-positive striatal neurons (**Fig. 3J**), suggesting reconstruction of nigrostriatal projections within the assembloids.

**Figure 3.**
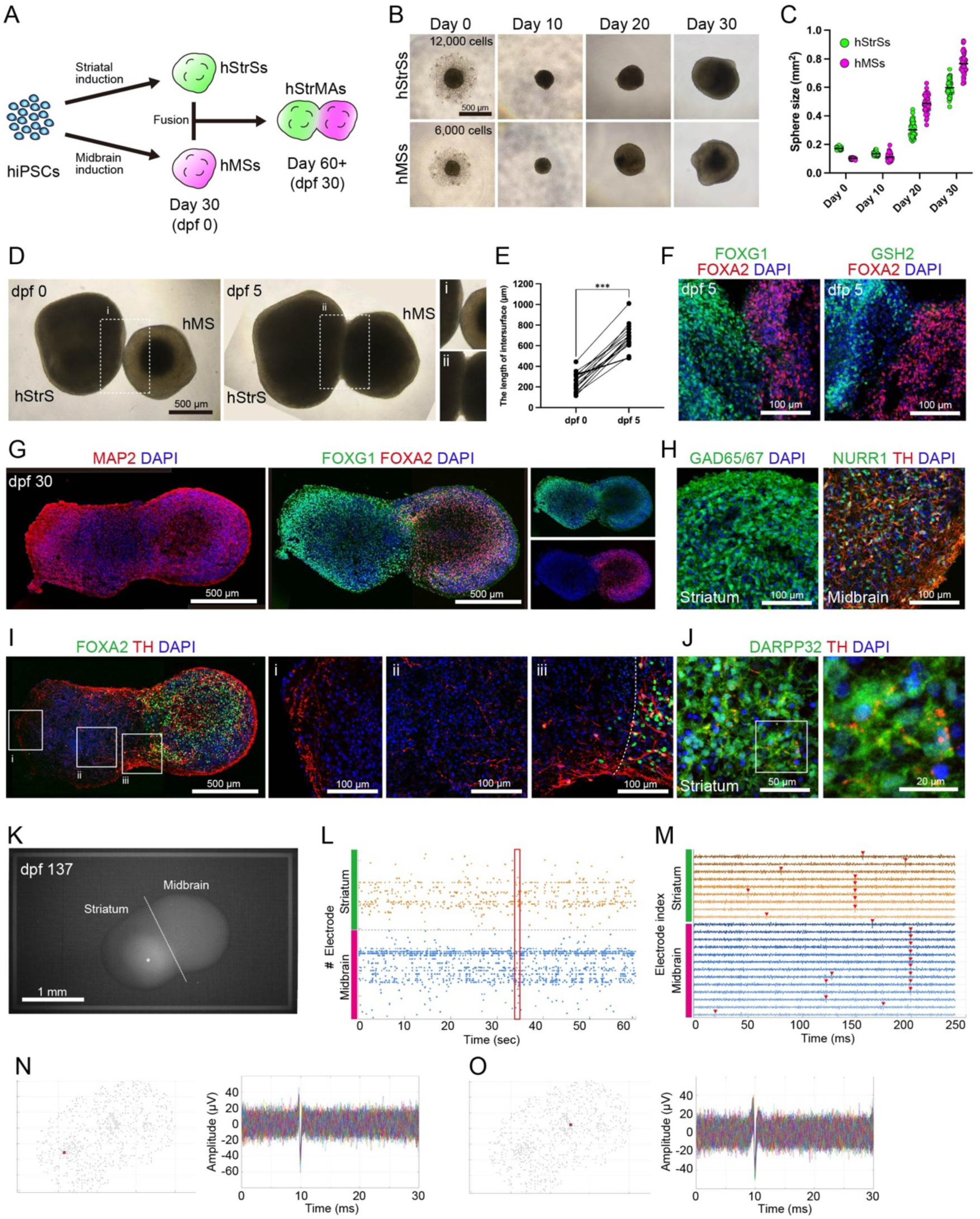
Functional neuronal projection in hStrMAs. (**A**) Schematic illustration of the experimental procedure for generating hStrMAs from hiPSCs. (**B**) Brightfield image of differentiating hStrSs (upper panels) and hMSs (lower panels) on days 0, 10, 20, and 30. (**C**) Quantification of spheroid size during differentiation (n = 43, mean ± SD). (**D**) Brightfield image before (0 dpf) and after (5 dpf) hStrS and hMS assembly. (**E**) Quantification of the intersurface length between 0 and 5 dpf. Significance (Student’s *t*-test): \*\*\**P* < 0.001 (n = 18, mean ± SD). (**F**) Immunostaining for FOXG1, FOXA2, and GSH2 at 5 dpf. (**G**) Immunostaining for MAP2, FOXG1, and FOXA2 at 30 dpf. (**H**) Immunostaining for GAD65/67 in the striatal region and NURR1 and TH in the midbrain region at 30 dpf. (**I**) Immunostaining for TH and FOXA2 at 30 dpf. (**J**) Immunostaining for DARPP32 and TH in the striatal region at 30 dpf. (**K**) Fluorescent image of an hStrMA on the HD-MEA at 137 dpf. Asterisk indicates autofluorescence. (**L**) Raster plots of the spike timing in hStrMAs detected at each electrode. (**M**) Neuronal spikes of selected channels that exhibited high-frequency neuronal activity. (**N**) Waveform of spontaneous neuronal spikes in the striatal region. (**O**) Waveform of spontaneous neuronal spikes in the midbrain region.

We then examined spontaneous neuronal activity in hStrMAs using a high-density multielectrode array (HD-MEA) system. To distinguish the striatum and midbrain after assembly, we generated hMSs from an EGFP-expressing hiPSC line obtained using the CRISPR-Cas9–based gene knock-in system (**Fig. S5**) and fused them with hStrSs generated from unlabeled hiPSCs (**Fig. 3K**). hStrMAs exhibited spontaneous neuronal activity in both the striatal and midbrain regions at 137 dpf (**Fig. 3L**). The action potentials detected from high active electrodes exhibited synchronized neuronal activities within each region (**Fig. 3M**). Representative waveforms of action potentials generated from striatal and midbrain neurons in the hStrMAs are presented in **Fig. 3N–O**.

To further investigate the cellular diversity of hStrMAs, we performed single-cell RNA sequencing (scRNA-seq) at 55 dpf (14,422 cells). Uniform manifold approximation and projection (UMAP) identified 15 distinct cellular clusters (**Figs. 4A and S6A–B**). Clusters were annotated according to the expression of representative regional and cell type-specific marker genes. Neuronal markers, including *MAP2*, *SYT1*, *MAPT*, and *MYT1L*, were broadly expressed across most clusters, indicating that most cells had acquired neuronal identity (**Figs. 4B and S6A, C**). Conversely, glial markers such as *S100B* and *SOX9* were detected in only 2.9% of all cells (**Fig. S6A**), indicating the clear segregation of regional identity at the single-cell level. *FOXG1* and *FOXA2* were expressed in mutually exclusive clusters (**Fig. 4C**). Striatal GABAergic neuron markers, including *GAD1*, *GAD2*, and *SLC32A1*, were enriched within forebrain-derived clusters. Contrarily, DA neuron markers such as *TH*, *DDC*, *SLC18A2*, and *SLC6A3* were expressed in VM clusters (**Figs. 4C and S6C**). Dot plots classified the composition of the striatal and VM cells based on region-specific gene expression profiles (**Fig. 4D**). Collectively, these results demonstrated that hStrMAs maintained distinct forebrain and VM neuronal populations at single-cell resolution.

**Figure 4.**
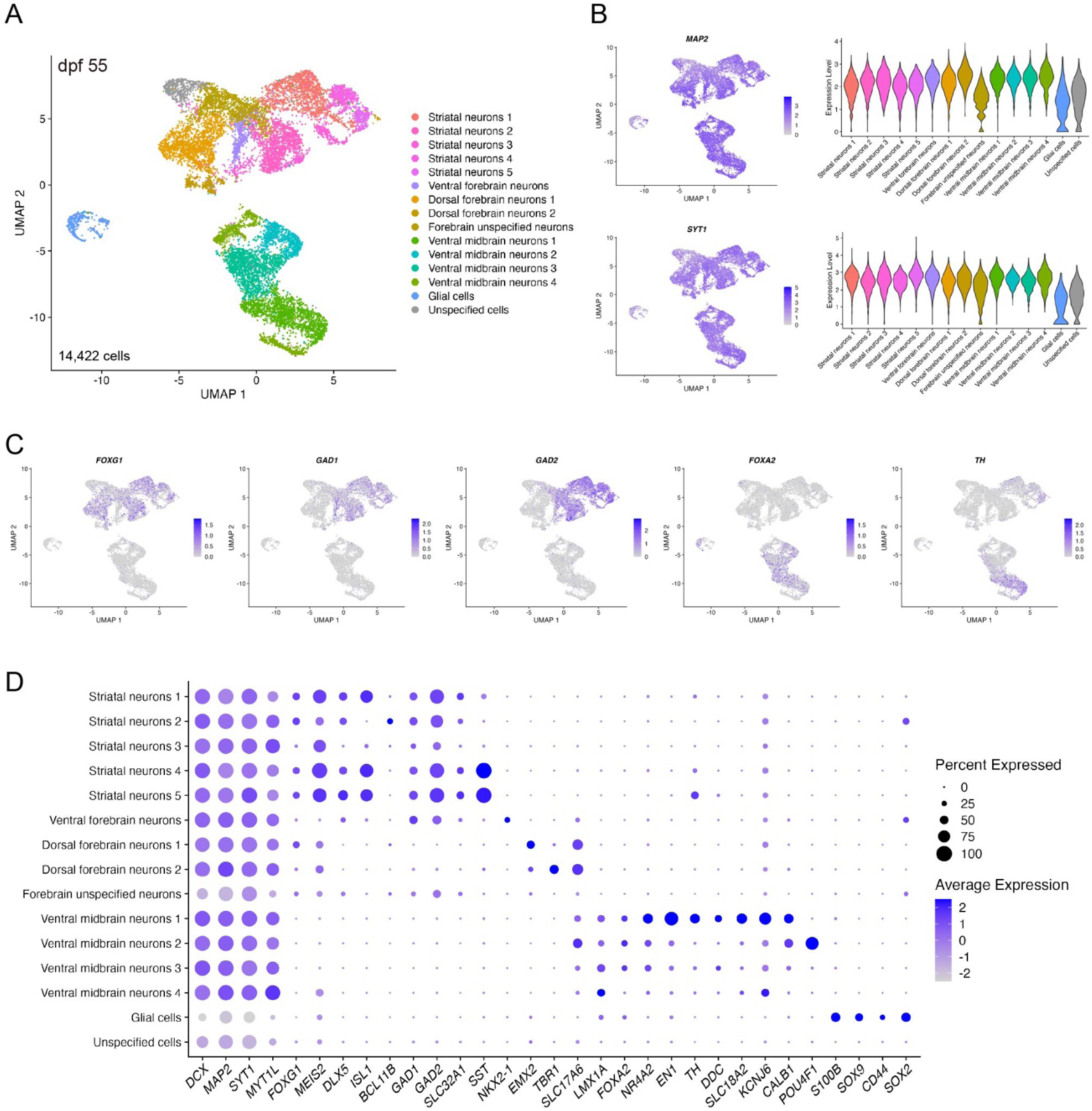
Characterization of hStrMAs using scRNA-seq. (**A**) UMAP of scRNA-seq data from 14,422 cells from hStrMAs at 55 dpf. (**B**) UMAP visualization and violin plots of *MAP2* and *SYT1.* (**C**) UMAP visualization of *FOXG1*, *GAD1*, *GAD2*, *FOXA2*, and *TH.* (**D**) Dot plots presenting the expression of representative genes in each cluster.

### Modeling synucleinopathy using hStrMAs

To model interregional α-synuclein propagation, we first established an experimental system that spatially and temporally controlled α-synuclein expression into hStrSs. We constructed a *piggyBac* vector to drive α-synuclein and mCherry expression under a DOX-responsive promotor (**Fig. S7A**) and then established a DOX-inducible α-synuclein–expressing hiPSC line. We confirmed that these cells normally expressed the pluripotent stem cell markers NANOG and OCT-3/4 (**Fig. S7B**) alongside α-synuclein and mCherry under DOX treatment, with immunocytochemistry and flow cytometry confirming that 94.5% of the cells expressed mCherry (**Fig. S7C–D**). Next, we induced hStrSs from the DOX-inducible α-synuclein–expressing hiPSCs, which were subsequently treated with 2 µg/mL DOX from day 45 of induction (**Fig. S7E**). The spheroid size of hStrSs did not differ between untreated and DOX-treated hStrSs (**Fig. S7F**), and mCherry fluorescence was maintained in DOX-treated hStrSs on day 75 (**Fig. S7G**). We also confirmed that α-synuclein was expressed in hStrSs, and the striatal neurons also expressed phosphorylated α-synuclein (**Fig. S7H**).

By restricting α-synuclein expression to the striatal region while eliminating endogenous α-synuclein expression in the midbrain region, we generated hStrMAs by fusing hStrSs induced from DOX-inducible α-synuclein–expressing hiPSCs and hMSs induced from *SNCA*-knockout hiPSCs (**Fig. S8**) (36). hStrMAs were treated with DOX for 30 days, and immunocytochemistry revealed that α-synuclein signals were detected in TH-positive DA neurons in the midbrain region.

Furthermore, we generated hStrMAs by fusing hStrSs induced from a DOX-inducible α-synuclein–expressing hiPSC line and hMSs induced from wild-type hiPSCs, confirming that mCherry and α-synuclein were expressed in the striatal region, and phosphorylated α-synuclein expression was induced by DOX treatment (**Fig. 5A**). In the midbrain region, α-synuclein was detected in TH-positive DA neurons, in which α-synuclein exhibited pathological features such as phosphorylation and fibrillization (**Fig. 5B**). Subsequently, we performed transcriptome analysis using bulk RNA-seq in the midbrain regions dissected from hStrMAs exposed to DOX for 30 days (**Fig. 5C**). Clustering analysis revealed a high correlation within each group (**Fig. 5D**). We found that DA neuron-related genes such as *TH*, *SLC18A2*, *SLC6A3*, *NR4A2*, *GCH1*, and *SNCA* were downregulated in the α-synuclein–propagated midbrain regions (**Fig. 5E**). GO analysis of genes downregulated in the α-synuclein–propagated midbrain regions revealed significant enrichment of cellular component terms related to synaptic function, including ‘postsynaptic membrane’ and ‘synaptic membrane’. In addition, biological process terms related to dopamine biosynthesis, such as ‘dopamine biosynthetic process’, ‘catechol-containing compound biosynthetic process’, and ‘catechol biosynthetic process’, were similarly enriched (**Fig. 5F**). Next, we highlighted individual genes contributing to these enriched GO terms in the α-synuclein–propagated midbrain regions (**Fig. 5G**). In addition to DA neuron-related genes, *TH*, *SLC6A3*, *SLC18A2*, *NR4A2*, and *GCH1*, the synaptic genes *SNCA*, *SV2B*, *ERC2*, and *APBA1* and the neural circuit-related genes *CHL1*, *DSCAM*, *NEGR1*, *KLHL1*, and *GPRIN3* were downregulated in the α-synuclein–propagated midbrain regions. These results indicated that α-synuclein propagation into the midbrain regions reduced DA and neural functioning.

**Figure 5.**
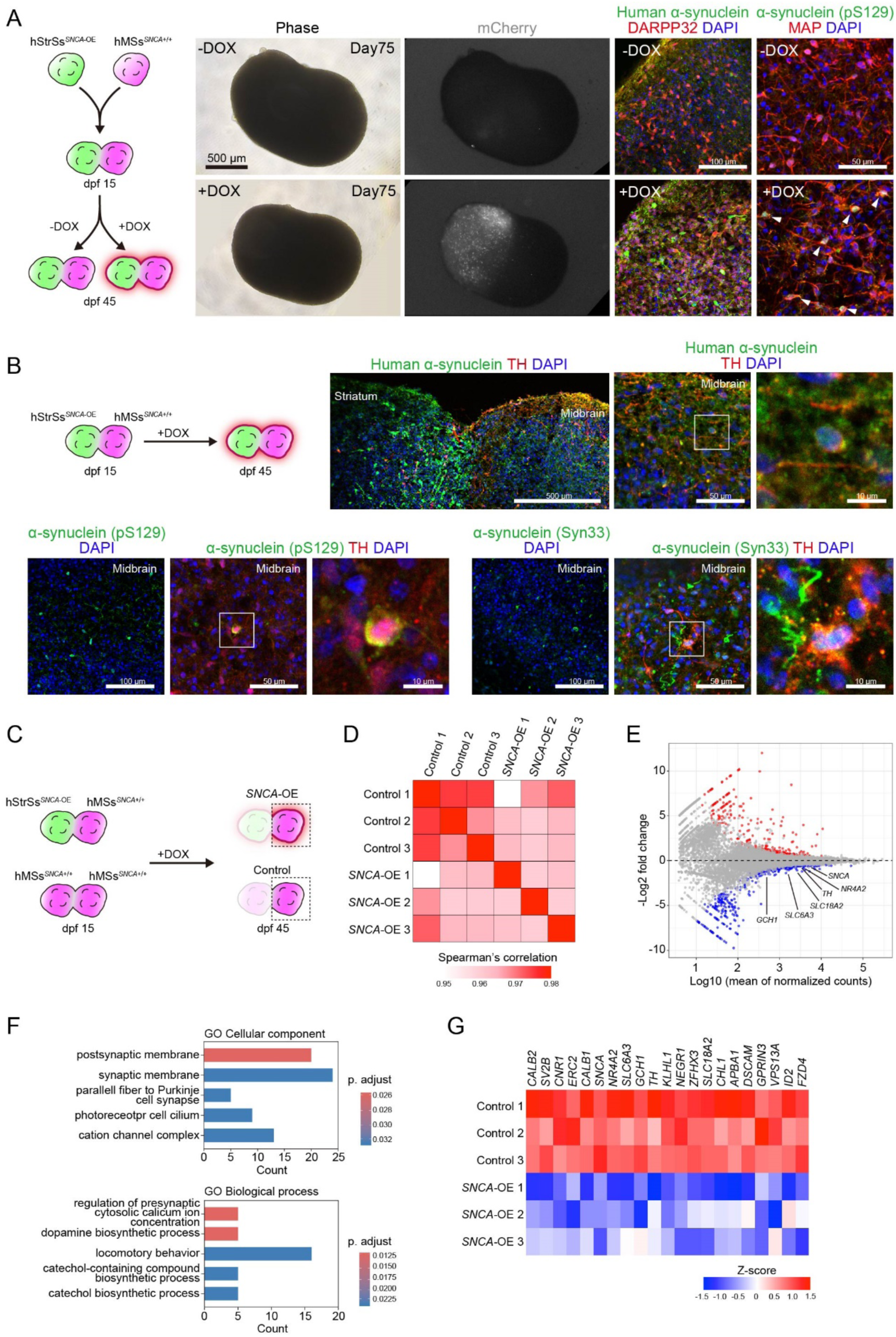
Modeling synucleinopathy in hStrMAs. (**A**) Induction of α-synuclein expression in the striatal region of hStrMAs. (**B**) Immunostaining for human α-synuclein, TH, α-synuclein (pS129), and α-synuclein (Syn33) in the α-synuclein–propagated hStrMAs. (**C**) Schematic illustration of the experimental procedure of RNA-seq analysis in hStrMAs. (**D**) Heatmap presenting Spearman’s correlation coefficients for α-synuclein–propagated midbrain regions and control midbrain regions. (**E**) MA plot of α-synuclein–propagated midbrain regions vs. control midbrain regions (adjusted p-value < 0.05, |log2 fold change| > 0.5). (**F**) Top five downregulated GO terms in α-synuclein–propagated midbrain regions. (**G**) presenting showing the genes highlighted in the GO terms.

## Discussion

The progressive distribution of α-synuclein pathology across brain regions is a pathological feature of PD, as reported in postmortem studies (9, 37). Although anatomical staging studies have suggested a stereotyped pattern of pathology progression, the mechanistic basis of interregional α-synuclein propagation in the human brain has remained difficult to dissect experimentally. Based on our previously established protocols, we constructed an hStrMA system in which α-synuclein expression can be spatially restricted and temporally induced after neural circuit formation, permitting investigation of the interregional α-synuclein propagation after the formation of nigrostriatal pathways.

Previous studies using animal models demonstrated that exogenously inoculated α-synuclein PFFs can induce seed aggregation and spreading pathology (12, 38). Similarly, 2D neuronal cultures allowed examination of the cell–cell transition of α-synuclein PFFs and the formation of Lewy body-like cytoplasmic inclusions and Lewy neurites (15, 39–41). Although these *in vivo* and *in vitro* models provided critical insights into templated misfolding and prion-like mechanisms (10), they do not fully recapitulate the nigrostriatal pathway in 3D human neural tissues.

Recent advances in human brain organoids and assembloids have permitted the reconstruction of region-specific neural tissues and interregional interactions *in vitro* (42–45). In addition, recent studies employed human midbrain organoid platforms to model α-synuclein–related pathology and vulnerability in PD (46–52). In addition, human neural assembloids have been used to reconstruct reciprocal neural network formation (26–28, 53) and investigate the early phenotypes of PD (26) and interregional α-synuclein transmission, aggregation, and dysfunction in DA neurons (29, 54). These studies collectively demonstrated the feasibility of modeling disease-relevant pathology within human 3D neural systems, representing important advances in the field.

In previously reported systems using either 3 × *SNCA* iPSCs or *SNCA*-overexpressing iPSCs, α-synuclein expression was already initiated during ongoing neural differentiation from undifferentiated iPSCs, making it difficult to separate pathology that was already initiated during neural induction from undifferentiated iPSCs from pathology that arises after neural circuit formation (29, 54). By contrast, our approach permits the spatially restricted and temporally controlled induction of α-synuclein after neural circuit formation, thereby experimentally defining the regional origin of pathology. This design allowed us to investigate whether synucleinopathy initiated in a specified brain region can propagate to a connected brain region after neural specification and maturation have already occurred. Thus, our experimental system can help clarify the molecular pathology of interregional α-synuclein propagation after the formation of nigrostriatal pathways within hStrMAs.

By selectively inducing α-synuclein in the striatal region after assembloid formation using a DOX-inducible system, our approach enabled the initiation of pathological protein expression after neural circuit formation within hStrMAs. Under conditions of temporally and spatially controlled α-synuclein expression, the ability of α-synuclein initiated in the striatal region to propagate to the midbrain region through an established neural network could be examined. The observed accumulation of α-synuclein pathology in the midbrain region supported the concept that the human neural circuit contributed to interregional propagation, consistent with anatomical observations in PD. Consequently, our assembloid system could represent a potential experimental model permitting separate investigation of the mechanisms of disease onset and progression after the formation of neural circuits, and it might be applicable to exploring novel therapeutic strategies.

Importantly, propagated α-synuclein in midbrain DA neurons exhibited pathological features, including phosphorylation and fibrillization, suggesting that neural circuit-mediated transfer is sufficient to trigger disease-relevant conformational conversion in recipient neurons. Transcriptomic analysis highlighted coordinated downregulation of DA identity genes (*TH*, *SLC6A3*, *SLC18A2*, *NR4A2*, and *GCH1*) in the midbrain region exposed to propagated α-synuclein. These genes constitute the core transcriptional network that maintains DA neuron identity and DA biosynthesis.

Although our model recapitulated the pathological features of synucleinopathy, several limitations remained. Despite pathological changes of α-synuclein in the midbrain region, further studies are required to determine the precise molecular mechanisms underlying transfer and the involvement of specific synaptic pathways. Although our assembloids reconstruct aspects of striatal–midbrain connectivity, the maturation and aging levels of our assembloids might be insufficient. Successful induction of aged assembloids will enhance the physiological relevance of the nigrostriatal pathway and permit examination of the phenotypes related to cellular senescence and aging. In addition, our assembloids lack a vascular system, immune cells, and matured glial cells, which can influence disease progression *in vivo*. Thus, the incorporation of glial cells or a vascular system might better recapitulate physiological environments at the multicellular level and permit exploration of various neurological diseases involving striatal–midbrain connections, such as Huntington’s disease (55), progressive supranuclear palsy (56) and multiple system atrophy (57).

In summary, we established an hStrMA system that enables spatially and temporally controlled α-synuclein induction and reveals interregional propagation. This controllable human experimental platform provides a foundation for dissecting the mechanisms of pathological protein spread and exploring therapeutic strategies targeting neural circuit-related aspects in neurodegenerative diseases.

## Materials and Methods

### Culture of hiPSCs

The hiPSC line 1231A3 was obtained from the Center for iPS Cell Research and Application, Kyoto University through the RIKEN Bioresource Research Center (Tsukuba, Japan) (58). The *SNCA* knockout (*SNCA*^−/−^) hiPSC line derived from 1231A3 cells was generated in a previous study (36). hiPSCs were cultured on iMatrix-511–coated six-well plates (Nippi, Tokyo, Japan) and maintained in Essential 8 medium or Essential 8 Flex medium (Thermo Fisher Scientific, Waltham, MA, USA) supplemented with 1% penicillin–streptomycin (Fujifilm Wako Pure Chemical Corporation, Osaka, Japan). For passaging, hiPSCs were dissociated into single cells using 0.5× TrypLE Select (Thermo Fisher Scientific) and replated at a density of 1,000–2,000 cells/cm^2^. The Rho-associate protein kinase inhibitor Y-27632 (10 μM; Selleck Chemicals, Houston, TX, USA) was added to the culture medium for the first 24 h after replating.

### Generation of a DOX-inducible *SNCA*-hiPSC line

Human *SNCA* cDNA was amplified using KOD One PCR Master Mix (TOYOBO Co. Ltd., Osaka, Japan) from cDNA samples of hiPSC-derived midbrain DA neurons and then cloned into the PB-TAC-ERP2 vector (#80,478, Addgene, Watertown, MA, USA) using InFusion HD (Takara Bio Inc., Kusatsu, Japan). Next, 1.0 × 10^5^ hiPSCs were transfected with 1.0 µg of the PB-TAC-ERP2-SNCA vector and 0.5 µg of the Super piggyBac Transposase Expression vector (System Biosciences, Palo Alto, CA, USA) using the Neon NxT Electroporation System (Thermo Fisher Scientific) under optimized conditions (1,200 V, 30 ms, 1 pulse) to. The cells were then plated onto an iMatrix-511–coated plate with Essential 8 medium supplemented with 10 µM Y-27632. Two days after electroporation, the medium was supplemented with 0.5 µg/mL puromycin (Takara Bio Inc.) for positive selection. Twelve clones were manually picked and expanded. Finally, we chose the appropriate clone displaying robust and stable mCherry expression. Target-specific primer sequences are listed in **Table S1**.

### Generation of an EGFP-expressing hiPSC line

The pRP[CRISPR]-hCas9-U6>AAVS1[gRNA#1] and pRP[Donor]-hAAVS1_LA:(EF1-EGFP-T2A-Puro-pA):hAAVS1_RA vectors, which were used to generate EGFP-expressing hiPSCs, were constructed and packaged by VectorBuilder Inc. (Chicago, IL, USA). The vector IDs are VB240508-1236dex and VB240504-1118jeq, respectively, which can be used to retrieve detailed information on vectorbuilder.com. hiPSCs (1.0 × 10^5^) were transfected with 0.5 µg of pRP[CRISPR]-hCas9-U6>AAVS1[gRNA#1] and 1.0 µg of pRP[Donor]-hAAVS1_LA:(EF1-EGFP-T2A-Puro-pA):hAAVS1_RA using the Neon NxT Electroporation System under optimized conditions (1,200 V, 30 ms, 1 plus). Two days after electroporation, the medium was supplemented with 0.5 µg/mL puromycin for positive selection. Target-specific primer sequences are listed in **Table S1**.

### Flow cytometry

The cells were dissociated into single cells using 0.5× TrypLE Select for 10 min at 37°C. The cells were stained with LIVE/DEAD Fixable Violet Dead Cell Stain Kit for excitation at 405 nm (Thermo Fisher Scientific) and then fixed with 4% paraformaldehyde (PFA) (Fujifilm Wako Pure Chemical Corporation) for 30 min at 4°C. The cells were suspended in PBS(−) and filtered through a nylon mesh (35-µm pore diameter) to eliminate large clumps before flow cytometry. Data were collected on a BD LSRFortessa flow cytometer (BD Biosciences, Franklin Lakes, NJ, USA) and analyzed using FlowJo software (BD Biosciences).

### Induction of hStrSs and hMSs from hiPSCs

Three-dimensional neurosphere formation was achieved using the SFEBq method as previously described (18). Briefly, hiPSCs were dissociated into single cells using 0.5× TrypLE Select and reaggregated by reseeding onto a low-cell attachment U-bottomed 96-well plate (Sumitomo Bakelite Co., Ltd., Tokyo, Japan) in Essential 8 medium supplemented with 10 μM Y-27632 (9,000 cells per well in 200 μL).

hStrSs were generated as previously described (30). Cells were differentiated in Essential 6 medium (Thermo Fisher Scientific) supplemented with 1% GlutaMax (Thermo Fisher Scientific), 1% nonessential amino acids (Fujifilm Wako Pure Chemical Corporation), 0.1 mM 2-mercaptoethanol (Fujifilm Wako Pure Chemical Corporation), and 1% penicillin–streptomycin. The differentiation medium was further supplemented with LDN193189 (200 nM, days 0–1; Stemgent, Beltsville, MD, USA), A83-01 (500 nM, days 0–6; Fujifilm Wako Pure Chemical Corporation), XAV939 (2 μM, days 0–11; Selleck Chemicals), and purmorphamine (0.5 μM, days 13–17; Selleck Chemicals). On days 5–11, the medium was gradually transitioned to AscleStem Neuronal Medium (Nacalai Tesque, Kyoto, Japan) supplemented with AscleStem Neuronal Supplement (Nacalai Tesque), 1% GlutaMax, and 1% penicillin–streptomycin. From day 17, neuronal maturation was induced using AscleStem Neuronal Medium supplemented with AscleStem Neuronal Supplement, 1% GlutaMax, and 1% penicillin–streptomycin, together with 10 μM DAPT (Selleck Chemicals), 20 ng/mL brain-derived neurotrophic factor (BDNF; PeproTech, Rocky Hill, NJ, USA), 200 μM ascorbic acid (Sigma-Aldrich, St. Louis, MO, USA), 1 μM PD0325901 (Selleck Chemicals), and 5 μM SU5402 (Selleck Chemicals). PD0325901 and SU5402 were withdrawn from the medium on day 24.

hMSs were generated according to previously described methods with minor modifications (31, 32). Cells were differentiated in Essential 6 medium supplemented with 1% GlutaMax, 1% nonessential amino acids, 0.1 mM 2-mercaptoethanol, and 1% penicillin–streptomycin. The medium was further supplemented with LDN193189 (200 nM, days 0–11), A83-01 (500 nM, days 0–6), purmorphamine (2 μM, days 1–16), CHIR99021 (1.5 μM, days 3–11; 7.5 μM, days 11–16; Selleck Chemicals), and FGF8B (100 ng/mL, days 9–16; PeproTech). On days 5–11, the medium was gradually transitioned from Essential 6 medium to AscleStem Neuronal Medium supplemented with AscleStem Neuronal Supplement, 1% GlutaMax, and 1% penicillin–streptomycin. From day 16, neuronal maturation was induced using AscleStem Neuronal Medium supplemented with AscleStem Neuronal Supplement, 1% GlutaMax, and 1% penicillin–streptomycin, together with 10 μM DAPT, 20 ng/mL BDNF, 200 μM ascorbic acid, 1 μM PD0325901 and 5 μM SU5402. From day 23, PD0325901 and SU5402 were withdrawn, and the medium was further supplemented with 10 ng/mL glial cell-derived neurotrophic factor (PeproTech), 500 μM dibutyryl-cAMP (Selleck Chemicals), and 2 ng/mL TGF-β3 (PeproTech). The differentiation medium was exchanged every other day.

### hStrMA generation

To generate hStrMAs, hStrSs and hMSs were generated from 12,000 and 6,000 cells per spheroid, respectively, and differentiated until day 16 (hMSs) or 17 (hStrSs) using the aforementioned protocols. The spheroids were subsequently maintained in AscleStem Neuronal Medium supplemented with AscleStem Neuronal Supplement, 1% GlutaMax, and 1% penicillin–streptomycin, together with 20 ng/mL BDNF and 200 μM ascorbic acid, until day 30. After 30 days of differentiation, hStrSs and hMSs were placed together into a single well of an ultralow attachment U-bottom 96-well plate to permit hStrMA formation. Two days after assembly, 10 μM DAPT, 1 μM PD0325901, and 5 μM SU5402 were added to the medium. From day 38, PD0325901 and SU5402 were removed from the medium.

### Immunocytochemistry

The hiPSC-derived cells, spheroids, and assembloids were fixed in 4% PFA for 30 min at 4°C. The spheroids and assembloids were transferred to 30% sucrose (Nacalai Tesque) solution and incubated overnight at 4°C. The samples were embedded within OCT compound (Sakura Finetek Japan Co., Ltd., Tokyo, Japan) and sectioned at 20 µm using a cryostat. The samples were then blocked and permeabilized in PBS containing 5% donkey serum (Jackson ImmunoResearch, West Grove, PA, USA) and 0.1% Triton X (Nacalai Tesque) for 60 min at room temperature and then incubated with primary antibodies (**Table S2**) overnight at 4°C. The samples were subsequently incubated with Alexa Fluor 488-, Alexa 555-, or Alexa Fluor 647-conjugated donkey secondary antibodies for 60 min at room temperature in the dark. The cell nuclei were counterstained with 2-(4-amidinophenyl)indole-6-carboxamidine dihydrochloride (1 μg/mL, Nacalai Tesque). Images were captured using a Leica TCS SPE laser confocal microscope (Leica Microsystems, Wetzlar, Germany).

### Fontana–Masson staining

hMSs were fixed in 4% PFA for 30 min at 4°C and washed with PBS. The samples were then stained using a Fontana–Masson Staining Kit (ScyTek Laboratories, West Logan UT, USA) according to the manufacturer’s protocol. Briefly, the samples were placed in ammoniacal silver solution and incubated in a 60°C water bath for 60 min until they became brown. The samples were washed with distilled water and then incubated with 0.2% gold chloride solution for 30 s. Subsequently, the samples were incubated with 5% sodium thiosulfate solution for 2 min.

### RNA extraction and quantitative PCR

Total RNA was extracted from spheroids using NucleoSpin RNA (Macherey-Nagel, Düren, Germany) and reverse-transcribed using a high-capacity cDNA reverse transcription kit with random hexamers according to the manufacturer’s protocol (Thermo Fisher Scientific). Quantitative PCR was performed using PowerUp SYBR green master mix (Thermo Fisher Scientific) on an ABI7500 real-time PCR system (Thermo Fisher Scientific). Data were analyzed using the ΔCT method with *GAPDH* expression as the internal control. Primer sequences are listed in **Table S1**.

### Bulk RNA-seq

Six spheroids were washed with PBS(−), quickly frozen, and stored at −80°C until use. RNA extraction and sequencing were outsourced to Cyberomix Inc. (Kyoto, Japan). Total RNA was extracted from neurospheres using the RNeasy Plus Micro Kit (Qiagen, Hilden, Germany) according to the manufacturer’s instructions. The total RNA concentration was measured using a Qubit 4 Fluorometer (Thermo Fisher Scientific), and samples with RNA integrity number > 7.0 were used for subsequent analysis. mRNA libraries were prepared using Universal Plus mRNA-seq with NuQuant kit (Tecan Group Ltd., Männedorf, Switzerland). The prepared libraries were sequenced on a DNBSEQ-G400RS system (MGI Tech, Shenzhen, China) in the paired-end mode (2 × 100 bp), generating approximately 33 million reads per sample (range, 20.6–43.7 million). Reads were aligned to the human reference genome (GRCh38) using HISAT2 (v2.2.1). Gene-level counts were obtained using featureCounts (v2.0.6). Differentially expressed genes (DEGs) were identified using DESeq2 (v1.36.0), and genes with adjusted p-value < 0.05 were considered significant (59). Transcriptomic data were visualized using R (RStudio, Boston, MA, USA), including pheatmap and ggplot2. Unbiased spatial mapping of the hStrSs and hMSs was performed using the VoxHunt algorithm (33).

### scRNA-seq

Three assembloids were washed with PBS(−) and then dissociated using 0.5× TrypLE Select at 37°C for 10 min. The dissociated cells were collected into 15-mL tubes with PBS(−) and centrifuged at 200 × *g* for 5 min. After removing the supernatant, the cells were resuspended in PBS(−) and filtered through a 20-µm cell strainer. Cells were counted and adjusted at a density of 1.05 × 10^5^ cells/200 µL. scRNA-seq was outsourced to Cyberomix Inc. The library was prepared using Chromium GEM-X Single Cell 3ʹ Reagent Kits v4 and the Chromium Controller according to the manufacturer’s instructions (10× Genomics, Pleasanton, CA, USA). The library was sequenced by NovaSeqX Plus (Illumina Inc., San Diego, CA, USA). From fastq files, quality control, alignment to the reference genome (GRCh38), and generation of count tables were performed using CellRanger v8.0.1 (10× Genomics). Further analyses were performed using the R package Seurat (v5.2.0). Cells with more than 500 or less than 6,000 detected genes or >10% mitochondrial content were also excluded.

### Patch–clamp analysis

The spheroids were embedded in 2% low melting point agarose XP (Nippon Gene Co., Ltd., Tokyo, Japan) and sectioned at 300 µm using a vibratome (VT1200S, Leica Microsystems) in ice-cold cutting solution bubbled with 95% O_2_/5% CO_2_. The cutting solution contained 2.5 mM KCl, 0.5 mM CaCl_2_, 10 mM MgSO_4_, 1.25 mM NaH_2_PO_4_, 3 mM sodium pyruvate, 92 mM N-methyl-D-glucamine, 20 mM HEPES, 12 mM N-acetyl-L-cysteine, 25 mM D-glucose, 5 mM L-ascorbic acid, and 30 mM NaHCO_3_. Slices were incubated at 34°C in a 1:1 mixture of cutting solution and artificial cerebrospinal fluid (ACSF; 125 mM NaCl, 3 mM KCl, 2 mM CaCl_2_, 1.3 mM MgCl_2_, 1.25 mM NaH_2_PO_4_, 10 mM D-glucose, and 25 mM NaHCO_3_) and then transferred to ACSF for subsequent recordings. Whole-cell patch–clamp recordings were performed from visually identified neurons in the spheroid slices under differential interference contrast optics using a Sutter patch–clamp amplifier (Sutter Instrument Company, Novato, CA, USA) under a microscope (BX-51WI, Olympus, Tokyo, Japan). Recording electrodes (4–8 MΩ) were pulled from borosilicate glass capillaries (GC150F-10, Harvard Apparatus, Holliston, MA, USA). The internal solution contained 130 mM potassium gluconate, 10 mM HEPES, 7 mM KCl, 0.2 mM EGTA, 4 mM NaCl, 2 mM MgATP, and 0.3 mM NaGTP, supplemented with 0.4% biocytin (pH 7.2, adjusted with KOH; 290–295 mOsm/kg). During recordings, slices were continuously perfused with ACSF at approximately 1.5 mL/min and maintained at approximately 29°C. Recordings with series resistance > 35 MΩ were excluded from the analysis. Capacitance was compensated, and bridge balance was adjusted during current–clamp recordings. Membrane potentials were not corrected for the liquid junction potential. Signals were low-pass–filtered at 5 kHz. Data were acquired and analyzed using Igor Pro (WaveMetrics, Lake Oswego, OR, USA) and SutterPatch software (Sutter Instrument Company, Novato, CA, USA).

### Calcium imaging analysis

hiPSC-derived neurons were transfected with an adeno-associated virus vector encoding the fluorescent calcium probe GCaMP6s (#100843-AAV1, Addgene) on day 40. The spontaneous activity of the hiPSC-derived neurons was recorded by fluorescence calcium imaging using the GCaMP6s probe. During recording, the cells were immersed in culture medium. Each recording was performed for 10 min at a frame rate of 20 Hz.

### HD-MEA analysis

A MaxOne HD-MEA chip (MaxWell Biosystems, Zürich, Switzerland) was used in this study. The electrodes were coated with 0.07% polyethyleneimine for 60 min and then with iMarix-511-silk for 60 min at 37°C. The assembloids were carefully transferred onto the electrodes, and the medium was removed using absorption spears (Agntho’s AB, Lidingö, Sweden), followed by 2 min of incubation. Then, 50 µL of the medium were gently added and incubated for 2 h. After assembloid attachment, 200 µL of the medium were gently added, followed by incubation overnight. On the next day, 800 µL of the medium were added to the chip. Half of the medium was exchanged every other day. All recordings were performed according to a previous report with minor modifications (60). Briefly, the spontaneous neuronal activity was recorded using MaxLab Live software (MaxWell Biosystems) at a sampling frequency of 20 kHz for 10 min. First, an activity scan assay was performed to identify the location of up to 1,020 active electrodes used for recording. Action potentials were bandpass-filtered with a frequency range of 300–3,000 Hz with a threshold of −5 × SD.

## Statistical analysis

The data were presented as the mean ± SD. The significance of differences was determined using Student’s *t*-test for single comparisons and by two-way ANOVA for multiple comparisons. Further statistical analysis for post hoc comparisons was performed using Tukey’s test. All statistical analyses were conducted using Prism 11 (GraphPad, Boston, MA, USA).

## Author contributions

K.N., N.A., and Y.M. designed research; K.N., N.A., Y.S., M.M., M.S., A.H., and M.N. performed research; K.N., N.A., Y.S., M.M., M.S., and A.H. contributed new reagents and analytic tools; K.N., N.A., Y.S., M.M., and M.S. analyzed data; K.N., N.A, Y.S., M.M., N.K., H.Y., A. H.-I., T.T., J.T., K.T., and Y.M. wrote the paper.

## Declaration of competing interest

The authors declare no conflict of interest.

## Acknowledgments

We thank Mrs. Sachiko Iwai for technical assistance and Dr. Hironobu Osaki for helpful discussion. This work was supported by the Grant-in-Aid for Scientific Research (C) (22K07382 to K.N.), Grant-in-Aid for Scientific Research (B) (26K03301 to Y.M.), Grant-in-Aid for Challenging Research (Exploratory) (24K22088 to Y.M.), Grant-in-Aid for Transformative Research Areas (A) (24H02332 to H.Y., 24H02333 to T.T., 24H02334 to A.H.-I., 24H02335 to Y.M., 25H02630 to M.M.) from the Ministry of Education, Culture, Sports, Science and Technology, Japan, the SDGs studies of Doshisha University (K.N.), the Cooperative Research Project Program of Tohoku University Research Institute of Electrical Communication (T.T. and H.Y.), the Kyoto Pharmaceutical University Fund for Collaborative Research (K.T.), the Uehara Memorial Foundation (K.N.), the Smoking Research Foundation (K.N.) and the Kobayashi Foundation (K.N.).

## Notes

### Competing Interest Statement

The authors have declared no competing interest.

