## Supplementary material for "Modeling interregional propagation of α-synuclein in human striatal–midbrain assembloids": Fig.S1-8, Table S1-2

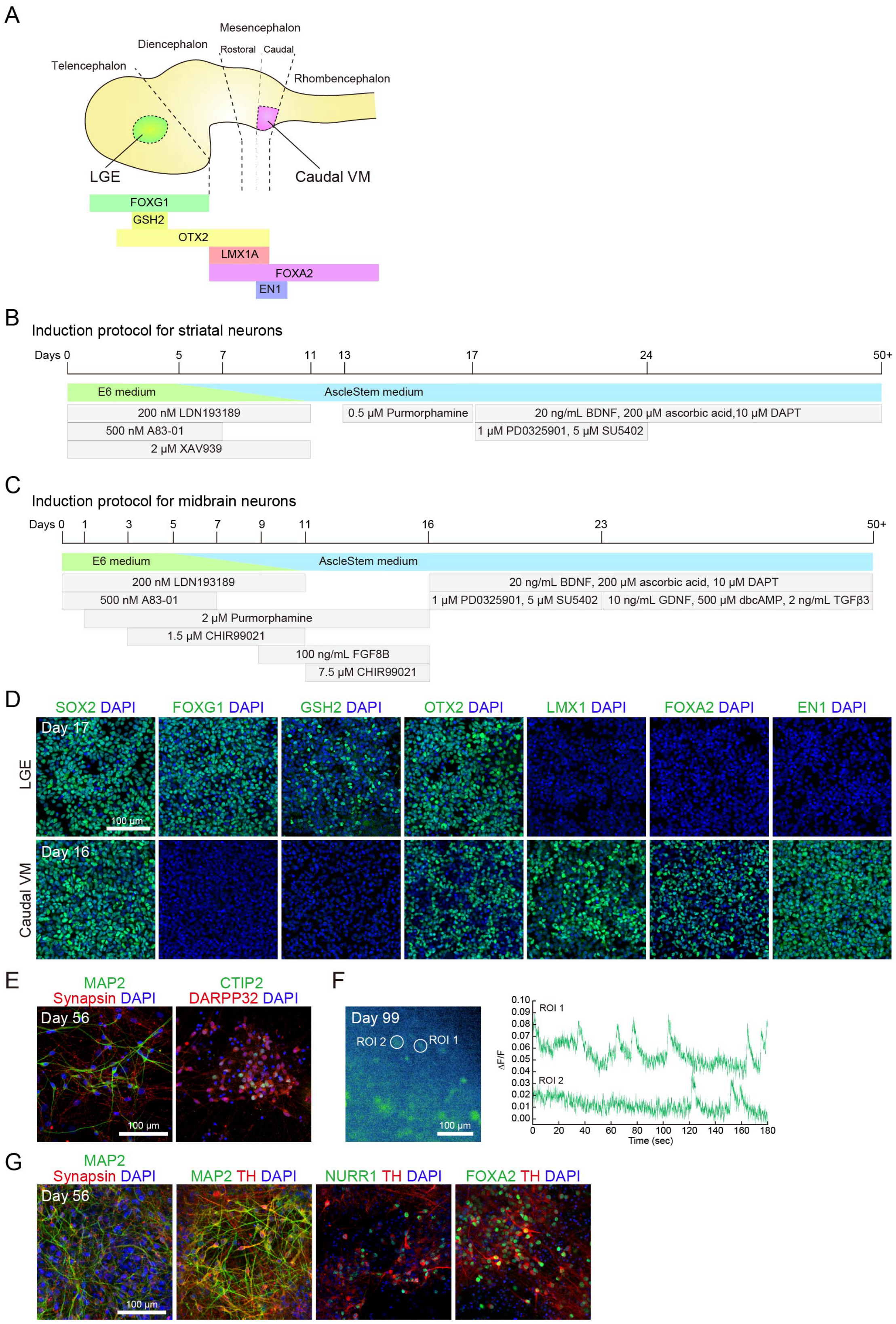


**Figure S1. Generation of striatal and midbrain neurons from hiPSCs in monolayer**

**culture.**

(**A**) Schematic illustration of the representative genes along the anterior–posterior axis. LGE, lateral ganglionic eminence; VM, ventral midbrain. (**B**) Protocol for generating striatal neurons from hiPSCs. BDNF, brain-derived neurotrophic factor. (**C**) Protocol for generating midbrain neurons from hiPSCs. FGF8B, fibroblast growth factor 8B; GDNF, glial cell-derived neurotrophic factor; TGFβ3, transforming growth factor-β3. (**D**) Immunostaining for SOX2, FOXG1, GSH2, OTX2, LMX1, FOXA2, and EN1 in LGE and caudal VM progenitors on days 17 and 16, respectively. (**E**) Immunostaining for MAP2, synapsin, CTIP2, and DARPP32 in striatal medium spiny neurons on day 56. (**F**) Calcium imaging analysis using GCaMP6s on day 99. ROI, region of interest. (**G**) Immunostaining for MAP2, synapsin, FOXA2, and TH in dopaminergic neurons on day 56.

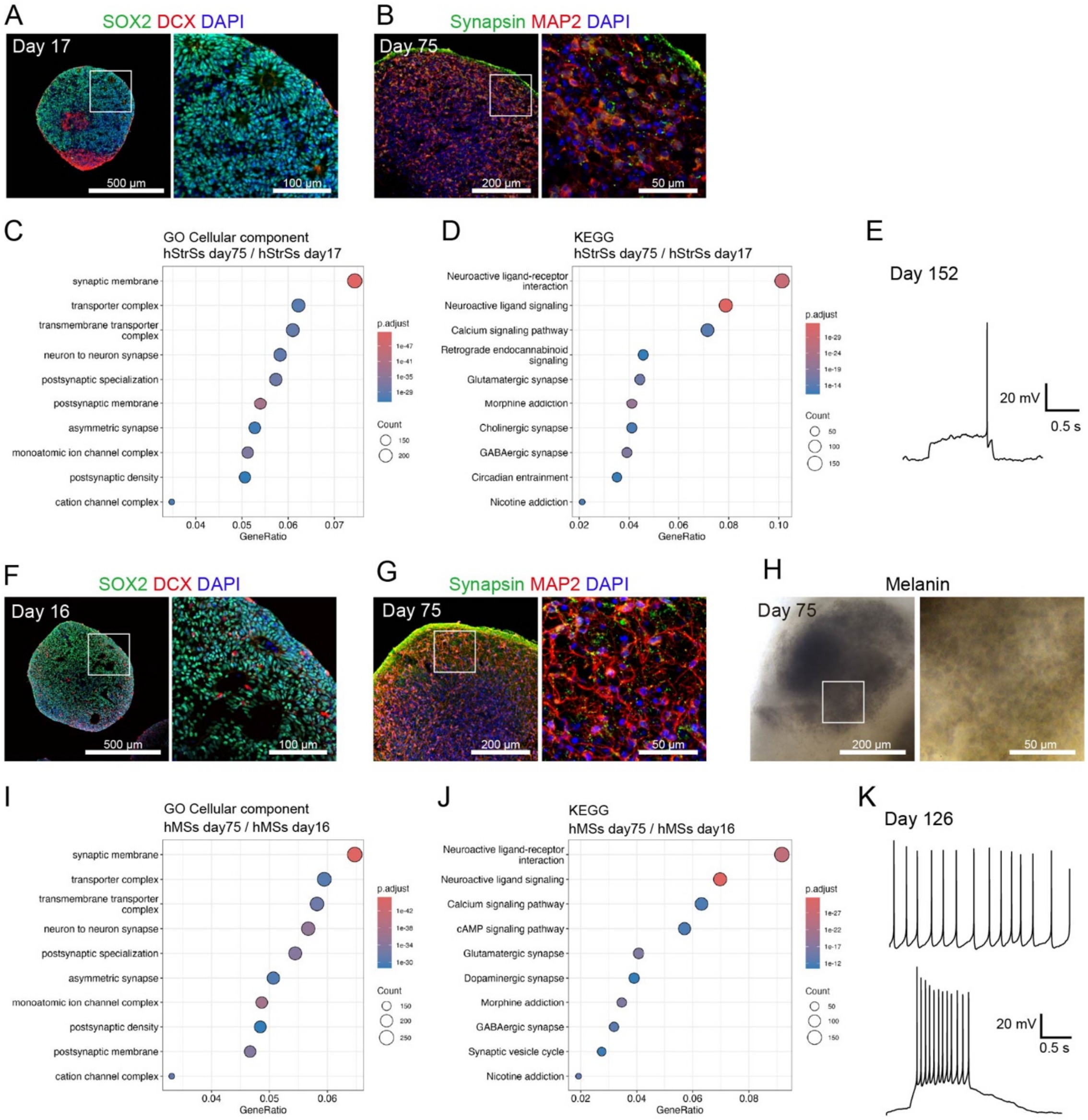


**Figure S2. Characterization of hStrSs and hMSs.**

(**A**) Immunostaining for SOX2 and DCX on day 17. (**B**) Immunostaining for synapsin and MAP2 on day 75. (**C**) Top 10 upregulated GO terms on day 75 versus day 17. (**D**) Upregulated KEGG pathways on day 75 versus day 17. (**E**) A representative late-spiking firing pattern during current–clamp recording on day 152 of culture. (**F**) Immunostaining for SOX2 and DCX on day 16. (**G**) Immunostaining for synapsin and MAP2 on day 75. (H) Fontana–Masson staining for melanin granules on day 75. (**I**) Top 10 upregulated GO terms on day 75 versus day 16. (**J**) Upregulated KEGG pathways on day 75 versus day 16. (**K**) Representative traces of spontaneous pacemaking activity (upper panel) and a firing pattern in response to a 4-pA depolarizing current injection during current–clamp recording (lower panel) after 126 days in culture.

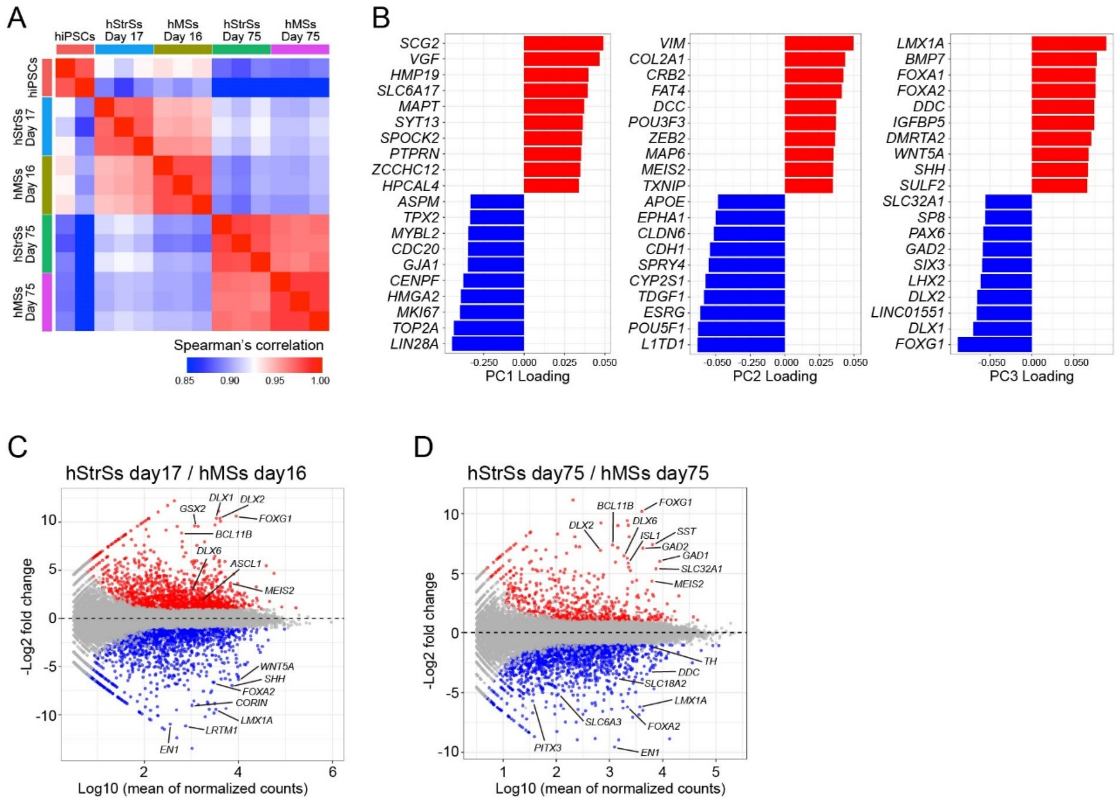


**Figure S3. Comparison of hStrSs and hMSs by bulk RNA sequencing.**

(**A**) Heatmap presenting Spearman's correlation coefficients among hiPSCs, hStrSs, and hMSs. (**B**) Genes with the highest loading on PC1 (left), PC2 (center), and PC3 (right). (**C**) MA plot presenting DEGs between hStrSs on day 17 and hMSs on day 16. Upregulated genes in hStrSs are presented in red, and downregulated genes in hStrSs are presented in blue (adjusted p-value < 0.05, |log2 fold change| > 1). (**D**) MA plot presenting DEGs between hStrSs and hMSs on day 75. Upregulated genes in hStrSs are presented in red, and downregulated genes in hStrSs are presented in blue (adjusted p-value < 0.05, |log2 fold change| > 1).

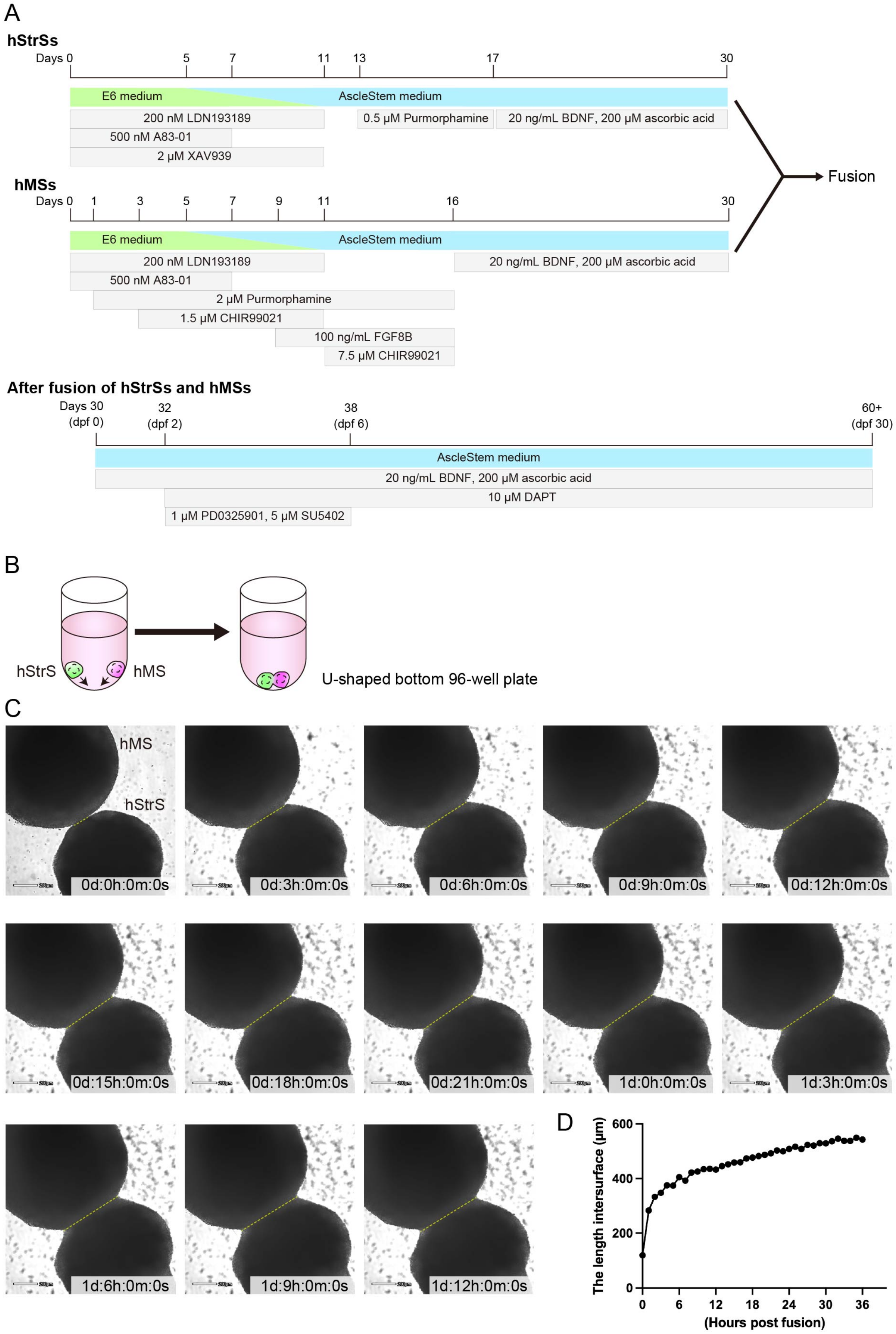
A
hStrSs
Days 0 5 7 11 13 17 30
E6 medium
AscleStem medium
200 nM LDN193189
500 nM A83-01
2 μM XAV939
0.5 μM Purmorphamine
20 ng/mL BDNF, 200 μM ascorbic acid
Fusion
hMSs
Days 0 1 3 5 7 9 11 16 30
E6 medium
AscleStem medium
200 nM LDN193189
500 nM A83-01
2 μM Purmorphamine
1.5 μM CHIR99021
100 ng/mL FGF8B
7.5 μM CHIR99021
20 ng/mL BDNF, 200 μM ascorbic acid
After fusion of hStrSs and hMSs
Days 30 (dpf 0)
32 (dpf 2)
38 (dpf 6)
60+ (dpf 30)
AscleStem medium
20 ng/mL BDNF, 200 μM ascorbic acid
10 μM DAPT
1 μM PD0325901, 5 μM SU5402
B
hStrS
hMS
U-shaped bottom 96-well plate
C
hMS
hStrS
0d:0h:0m:0s
0d:3h:0m:0s
0d:6h:0m:0s
0d:9h:0m:0s
0d:12h:0m:0s
0d:15h:0m:0s
0d:18h:0m:0s
0d:21h:0m:0s
1d:0h:0m:0s
1d:3h:0m:0s
1d:6h:0m:0s
1d:9h:0m:0s
1d:12h:0m:0s
D
The length intersurface (μm)
600
400
200
0
0 6 12 18 24 30 36
(Hours post fusion)

**Figure S4. Generation and assembly of hStrSs and hMSs.**

(**A**) Protocol for generating hStrMAs from hiPSCs. BDNF, brain-derived neurotrophic factor; FGF8B, fibroblast growth factor 8B. (**B**) Schematic illustration of the assembly of hStrSs and hMSs. (**C**) Time-lapse imaging during hStrS and hMS assembly. (**D**) Quantification of the intersurface length during hStrS and hMS assembly.

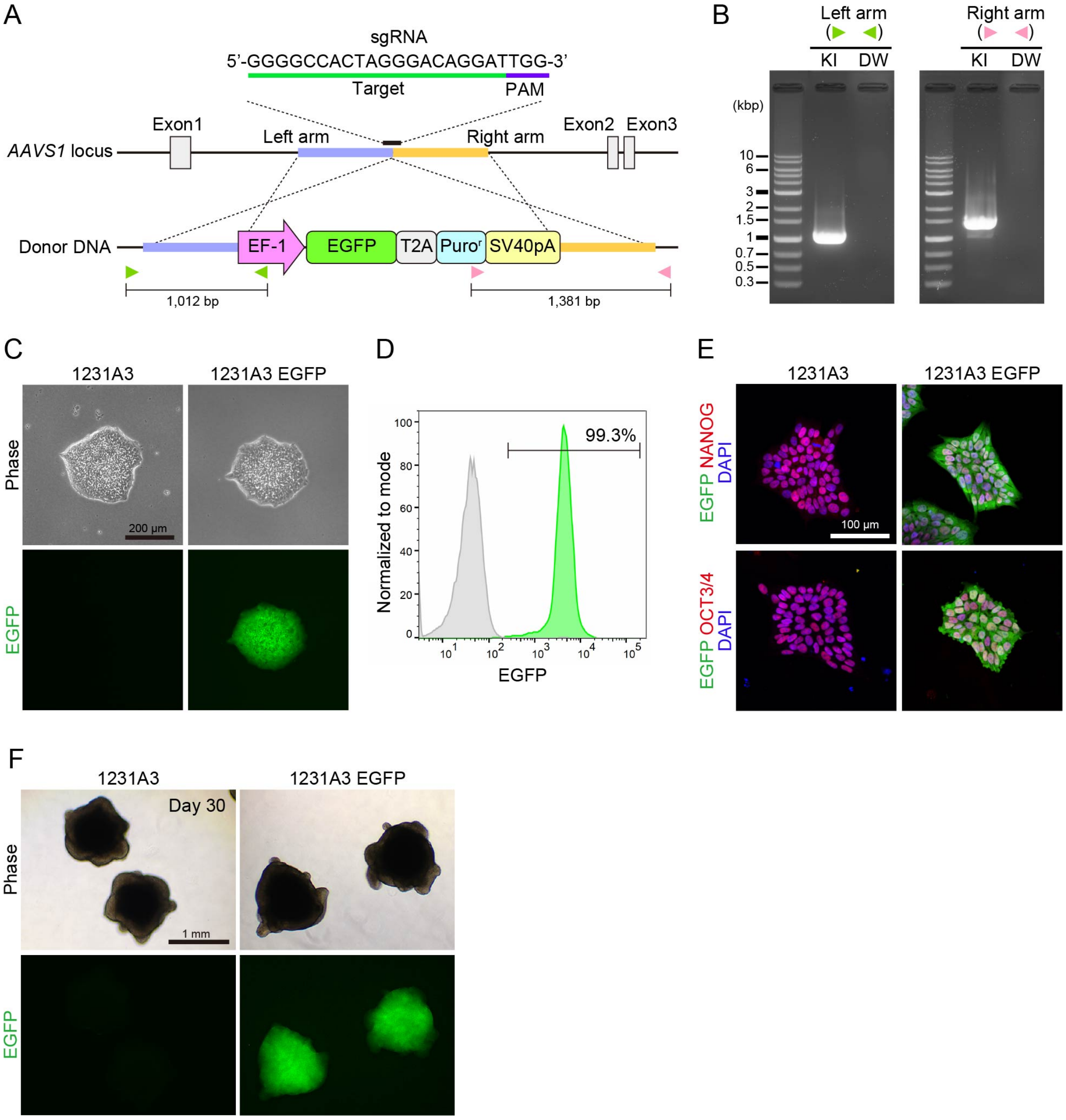


**Figure S5. Generation of hiPSCs with *EGFP* knock-in.**

(**A**) Schematic illustration of *EGFP* knock-in using the CRISPR-Cas9 system into the AAVS1 locus. (**B**) Genotyping of the *EGFP*–knocked-in hiPSC line. (**C**) Phase contrast and EGFP fluorescence imaging of the *EGFP*-knocked-in hiPSC line. (**D**) Flow cytometry of EGFP fluorescence in the *EGFP*-knocked-in hiPSC line. (**E**) Immunostaining of GFP, NANOG, and OCT3/4 in the *EGFP*-knocked-in hiPSC line. (**F**)

Phase contrast and EGFP fluorescence imaging of hMSs generated from the *EGFP*-knocked-in hiPSC line.

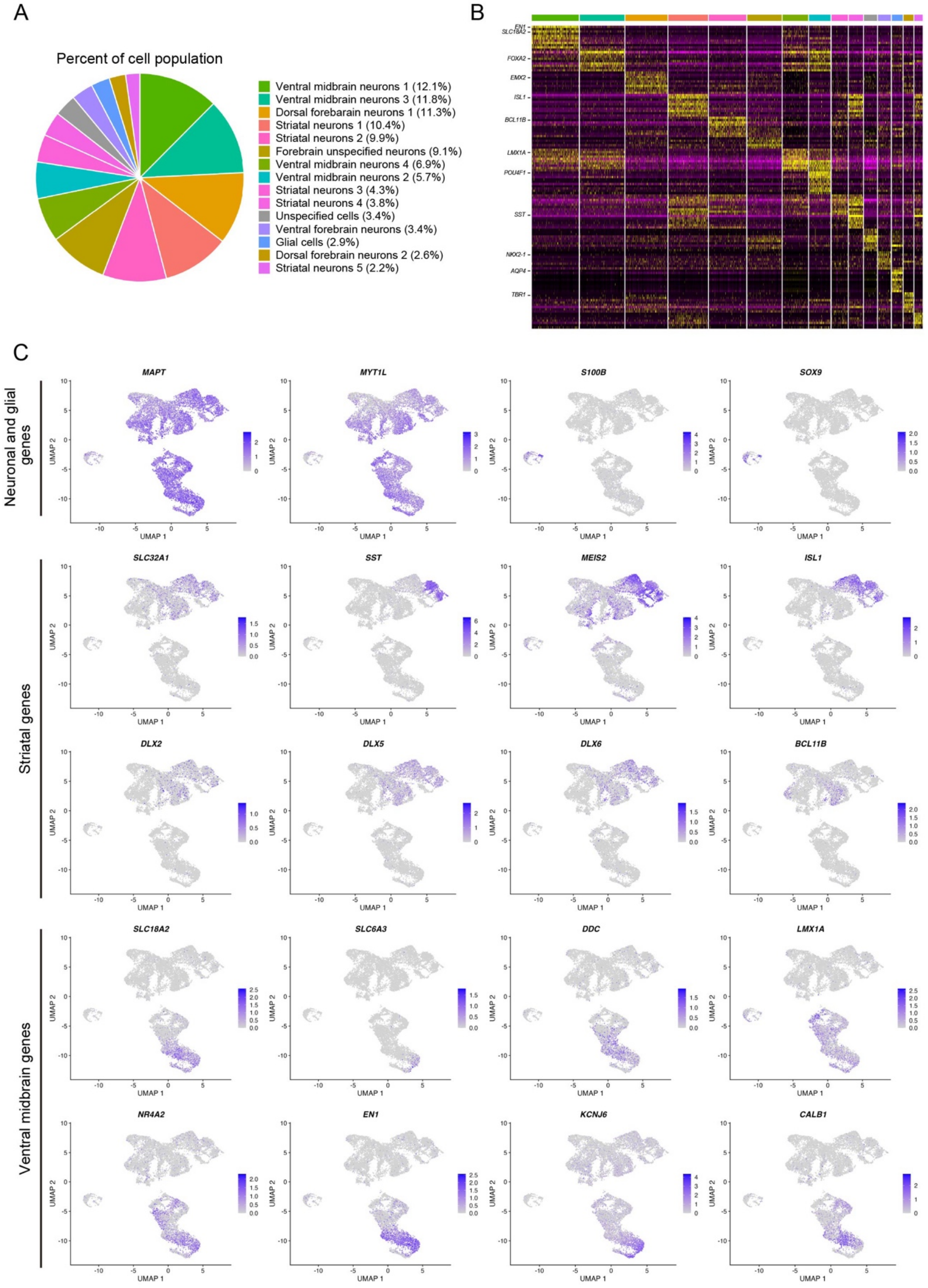
A
Percent of cell population
Ventral midbrain neurons 1 (12.1%)
Ventral midbrain neurons 3 (11.8%)
Dorsal forebarain neurons 1 (11.3%)
Striatal neurons 1 (10.4%)
Striatal neurons 2 (9.9%)
Forebrain unspecified neurons (9.1%)
Ventral midbrain neurons 4 (6.9%)
Ventral midbrain neurons 2 (5.7%)
Striatal neurons 3 (4.3%)
Striatal neurons 4 (3.8%)
Unspecified cells (3.4%)
Ventral forebrain neurons (3.4%)
Glial cells (2.9%)
Dorsal forebrain neurons 2 (2.6%)
Striatal neurons 5 (2.2%)
B
EN1
SLC18A2
FOXA2
EMX2
ISL1
BCL11B
LMX1A
POU4F1
SST
NKX2-1
AQP4
TBR1
C
Neuronal and glial genes
MAPT
MYT1L
S100B
SOX9
Striatal genes
SLC32A1
SST
MEIS2
ISL1
DLX2
DLX5
DLX6
BCL11B
Ventral midbrain genes
SLC18A2
SLC6A3
DDC
LMX1A
NR4A2
EN1
KCNJ6
CALB1
UMAP 1
UMAP 2

**Figure S6. Single-cell RNA sequencing of hStrMAs.**

(**A**) Percentage of each cluster in hStrMAs. (**B**) Heatmap for the top 10 genes in each cluster. (**C**) UMAP visualization presenting the representative clusters of neuronal, striatal, and ventral midbrain genes.

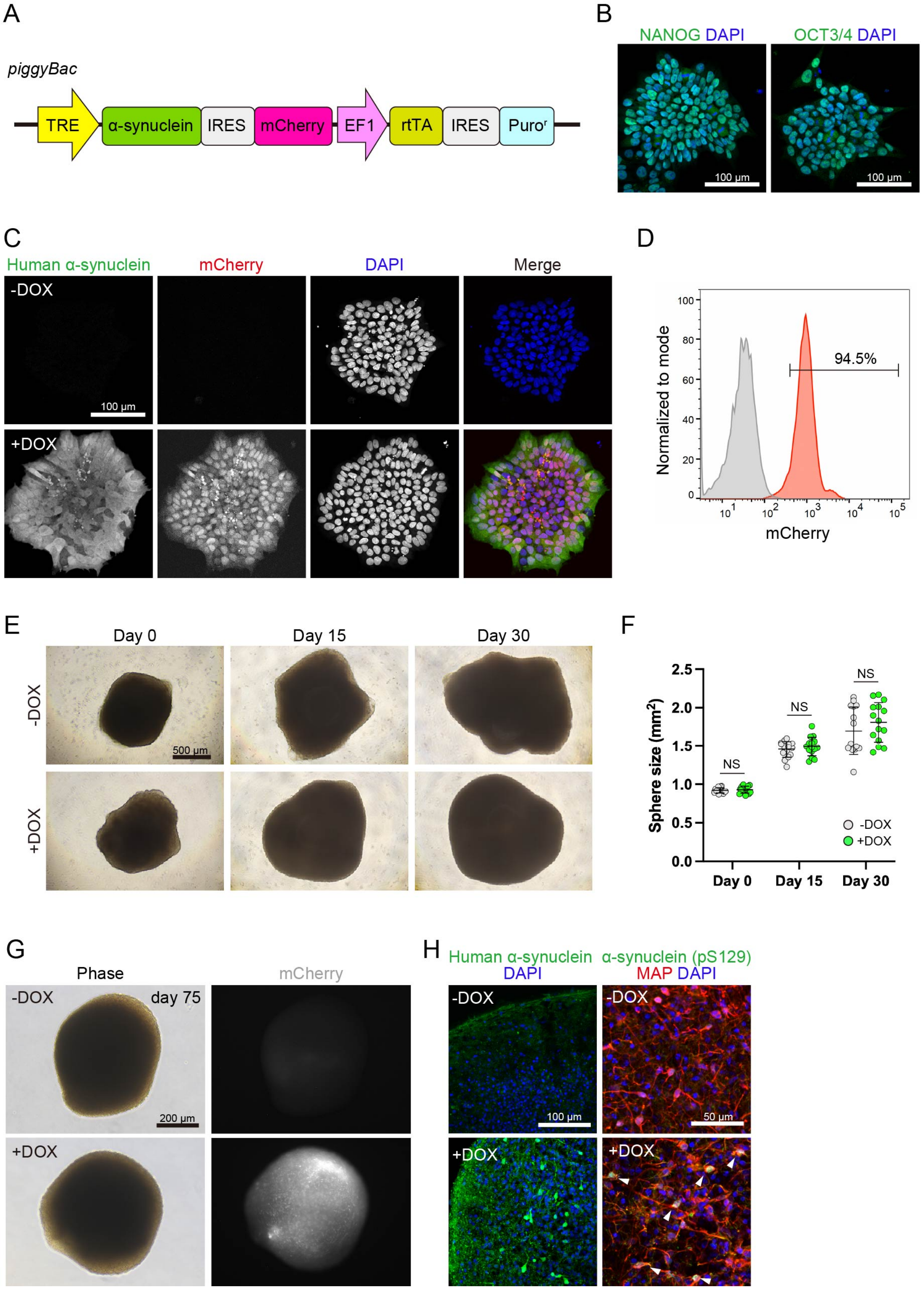

A
piggyBac
TRE
α-synuclein
IRES
mCherry
EF1
rtTA
IRES
Puro$^r$
B
NANOG DAPI
OCT3/4 DAPI
100 μm
100 μm
C
Human α-synuclein
mCherry
DAPI
Merge
-DOX
+DOX
100 μm
D
Normalized to mode
94.5%
mCherry
E
Day 0
Day 15
Day 30
-DOX
+DOX
500 μm
F
Sphere size (mm$^2$)
NS
NS
NS
-DOX
+DOX
Day 0
Day 15
Day 30
G
Phase
mCherry
-DOX
day 75
+DOX
200 μm
H
Human α-synuclein DAPI
α-synuclein (pS129) MAP DAPI
-DOX
+DOX
100 μm
50 μm

**Figure S7. Characterization of DOX-inducible α-synuclein–expressing hiPSCs.**

(**A**) Map of the DOX-inducible α-synuclein–expressing *piggyBac* vector. (**B**) Immunostaining for NANOG and OCT3/4 in DOX-inducible α-synuclein–expressing hiPSCs. (**C**) Immunostaining for human α-synuclein and mCherry fluorescence after DOX treatment. (**D**) Flow cytometry of mCherry fluorescence after DOX treatment. (**E**) Brightfield image during DOX treatment on days 0, 15, and 30. (**F**) Quantification of spheroid size during DOX treatment (n = 15, mean ± SD). NS, not significant. (**G**) Brightfield image and mCherry fluorescence of DOX-treated hStrSs. (**H**) Immunostaining for human α-synuclein, α-synuclein (pS129), and MAP2 in DOX-treated hStrSs.

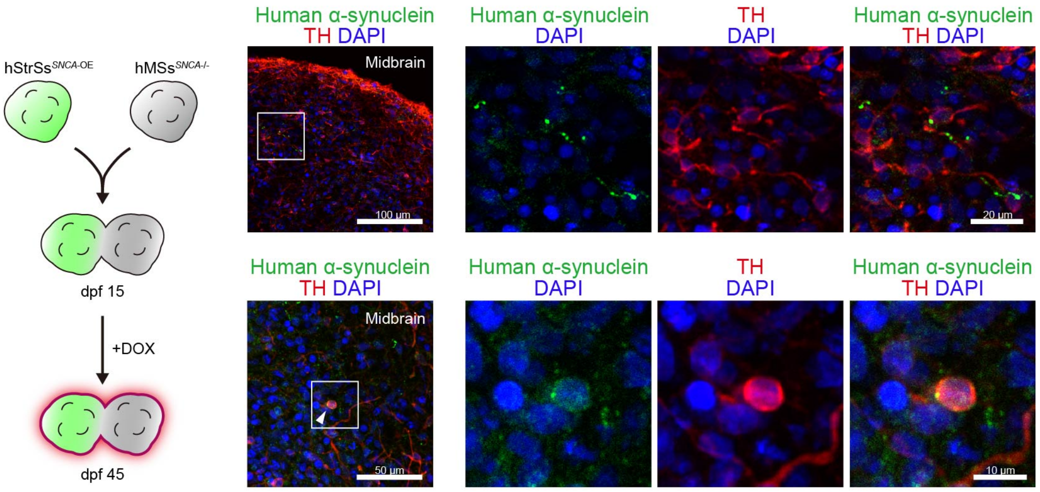


**Figure S8. Characterization α-synuclein propagation across brain regions.**

Immunostaining of human α-synuclein and TH in the midbrain region of hStrMAs.

### Table S1. List of primers

| Primers | Target | Primer sequence (5′-3′) |
|---|---|---|
| Cloning | *SNCA* | Fw: GCGGTGGCGGCCATCATGGATGTATTCATGAAAGG |
| | (423 bp) | Rv: CGTTTTAGCTAGATCTTAGGCTTCAGGTTCGTAGT |
| Genotyping | AAVS1 LA | Fw: CACTGTTTCCCCTTCCCAGG |
| | (1,012 bp) | Rv: ACTGCACTTATATACGGTTC |
| | AAVS1 RA | Fw: TTCTACGAGCGGCTCGGCTT |
| | (1,381 bp) | Rv: GAAGAGTGAGTTTGCCAAGCAG |
| qPCR | *FOXA2* | Fw: TTCAGGCCCGGCTAACTCT |
| | (67 bp) | Rv: AGTCTCGACCCCCACTTGCT |
| | *GAPDH* | Fw: TTGAGGTCAATGAAGGGGTC |
| | (117 bp) | Rv: GAAGGTGAAGGTCGGAGTCA |
| | *GSH2* | Fw: ATTCCACTGCCTCACCATGG |
| | (91 bp) | Rv: CAGGAGTTGCGTGCTAGTGA |
| | *LMX1A* | Fw: GATCCCTTCCGACAGGGTCTC |
| | (175 bp) | Rv: GGTTTCCCACTCTGGACTGC |

### Table S2. List of antibodies

| Antibody | Host | Dilution | Source | Catalog # |
|---|---|---|---|---|
| α-Synuclein (pS129) | Mouse | 500 | WAKO | 015-25191 |
| α-Synuclein (Syn33) | Rabbit | 500 | Merck Millipore | ABN2265 |
| CTIP2 | Rat | 2,000 | Abcam | ab8465 |
| DARPP32 | Rabbit | 200 | Abcam | ab40801 |
| DARPP32 | Mouse | 500 | Santa Cruz | sc-271111 |
| DCX | Mouse | 500 | Santa Cruz | sc-271390 |
| EN1 | Rabbit | 600 | GeneTex | GTX81554 |
| FOXA2 | Goat | 500 | R&D Systems | AF2400 |
| FOXG1 | Rabbit | 500 | Abcam | ab18259 |
| GAD65/67 | Rabbit | 500 | SIGMA | G5163 |
| GFP | Rabbit | 500 | MBL | 598 |
| GSH2 | Rabbit | 500 | Merck Millipore | ABN162 |
| Human α-synuclein | Mouse | 1,000 | Santa Cruz | sc-58480 |
| LMX1 | Rabbit | 3,000 | Merck Millipore | AB10533 |
| MAP2 | Mouse | 1,000 | Sigma-Aldrich | M4403 |
| MAP2 | Rabbit | 500 | Merck Millipore | AB5622 |
| NANOG | Goat | 40 | R&D Systems | AF1997 |
| NURR1 | Mouse | 500 | Perseus Proteomics | PP-N1404-00 |
| OCT3/4 | Mouse | 200 | Santa Cruz | sc-5279 |
| OTX2 | Goat | 500 | R&D Systems | AF1979 |
| SOX2 | Rabbit | 500 | Merck Millipore | AB5603 |
| Synapsin | Rabbit | 1,000 | Merck Millipore | 574,777 |
| TAU | Mouse | 1,000 | Merck Millipore | MAB3420 |
| TH | Rabbit | 500 | Merck Millipore | AB152 |
| TH | Mouse | 500 | Sigma-Aldrich | T2928 |

**Movie S1.** Brightfield image of the fusion of a hStrS and hMS.
